# Calorie Restriction Prevents Age-Related Changes in the Intestinal Microbiota

**DOI:** 10.1101/2020.09.02.279778

**Authors:** Kavitha Kurup, Stephanie Matyi, Cory B. Giles, Jonathan D. Wren, Kenneth Jones, Aaron Ericsson, Daniel Raftery, Lu Wang, Daniel Promislow, Arlan Richardson, Archana Unnikrishnan

## Abstract

The effect of calorie restriction (CR) on the microbiome, fecal metabolome, and colon transcriptome of adult and old male mice was compared. Life-long CR increased microbial diversity and the *Bacteriodetes*/*Fermicutes* ratio and prevented the age-related changes in the microbiota, shifting it to a younger microbial and fecal metabolite profile in both C57BL/6JN and B6D2F1 mice. Old mice fed CR were enriched in the *Rikenellaceae, S24-7* and *Bacteroides* families. The changes in the microbiome that occur with age and CR were initiated in the cecum and further modified in the colon. Short-term CR in adult mice had a minor effect on the microbiome but a major effect on the transcriptome of the colon mucosa. These data suggest that the primary impact of CR is on the physiological status of the gastrointestinal system, maintaining it in a more youthful state, which in turn results in a more diverse and youthful microbiome.

## Introduction

The first and the most studied manipulation shown to increase lifespan in mammals is caloric restriction (CR). The classic study by McCay et al (1935) showed that one could increase the lifespan of rats by dramatically reducing their food consumption. Since this initial observation, numerous laboratories have confirmed these results and have shown that reducing food consumption 30 to 50% (without malnutrition) consistently increases both the mean and maximum lifespans of both laboratory rats and mice (Weindruch and Walford, 1988; Masoro, 2005). For example, Turturro et al (1999) showed that 40% CR increased the lifespan of four inbred strains of male and female laboratory mice and three inbred strains of male and female laboratory rats. The effect of CR on longevity is not limited to rodents as CR has been shown to increase the lifespan of a large number of diverse animal models ranging from invertebrates (yeast, *C*. *elgans*, and *Drosophila*) to dogs and non-human primates (Unnikrishnan et al., 2019). Because CR has a broad effect on lifespan, it is generally considered that the effect of CR on lifespan is universal, i.e., it occurs in all organisms.

Because the gastrointestinal (GI) system is the first organ/tissue that encounters the impact of reduced food consumption, there have been several studies on the effect of CR on the GI-system. Early studies, primarily from Peter Holt’s laboratory, showed that CR had a major impact on the colon. CR was shown to prevent/delay the age-related changes in rat colon, e.g., crypt hyperplasia, and expression of mucosal enzymes (Heller et al., 1990; Holt et al., 1991), reduced cell proliferation (Steinbach et al., 1993), and enhance apoptosis (Holt et al., 1998). As expected from these findings, CR has also been shown to reduce colorectal cancer in various rodent models (Klurfeld et al., 1987; Kritchevsky, 1993; Olivo-Marston et al., 2014). With the advent of 16S rRNA sequencing and metagenomics, it is now possible to interrogate the colon microbiome and study the effect of CR. Two groups have reported that long-term CR had a significant impact on the microbiome of old mice (Zhang et al., 2013; Kok et al., 2018). These studies were conducted with aging colonies of mice specific to those particular laboratories and for which there were no lifespan data. Because the institutional animal husbandry environment and the health status of the host can have a major impact on the microbiome (Ericsson et al., 2018), we felt it was important to establish the effect of CR on the microbiome of well characterized mice from the aging colony maintained by National Institute on Aging (NIA). Out data show that life-long CR had a major impact on the colon microbiome of both C57BL/6JN and B6D2F1 mice, resulting in a ‘younger’ appearing microbiome and these changes resulted from the effect of CR on the gastrointestinal system.

## Results

### Effect of age and caloric restriction on the gut microbiome of mice

Using male mice obtained from the aging colony maintained by the NIA, we studied the microbiota composition of the cecum and colon of 9- and 24-month old male C57BL/6JN mice fed AL or CR (started at 14 weeks of age) by sequencing the V4 region of the bacterial 16S rRNA gene. These data are presented in Tables 1S and 2S in the supplement. Because genotype of the host can impact the microbiome, we also analyzed the microbiome of male B6D2F1 mice obtained from the NIA. Adult B6D2F1 were not available from the NIA; therefore, we were only able to measure the microbiome of the cecum and colon of 24-month-old B6D2F1 mice fed either AL or CR, and these data are presented in Table 3S in the supplement. The data in Tables 1S, 2S, and 3S were analyzed by principal component analysis (PCA), and the results are shown in Figure 1. For C57BL/6JN mice (Figure 1A and B), adult mice fed AL or CR and the old mice fed CR grouped together for both the cecum and colon. Importantly, old AL mice were clearly segregated from the other three groups for both the cecum and colon, indicating that CR attenuated the effect of age on the microbiome in the old mice. In old B6D2F1 mice (Figure 1C and D), we observed a clear separation in the colon microbiome for mice fed AL and CR; however, the microbiome of the cecum of the AL and CR groups overlapped primarily because of one animal (Figure 1C). Figure 1 also shows that when projecting all of the data in the same space, the old mice fed AL consistently occurred on the left side of the plots, indicating that age and CR have similar metagenomic effects in the two strains of mice in both the cecum and colon. In Figure 1E and F, we directly compared the microbiome data for the cecum or colon in old C57BL/6JN and B6D2F1 mice. In the cecum, there was very little separation between the four groups. In contrast, we observed a clear separation between AL and CR mice in the colon, suggesting that the effect of CR on the microbiome becomes modified as the fecal material transits to the colon. Interestingly, the C57BL/6JN and B6D2F1 mice grouped together for both AL and CR, indicating that diet was a more important variable than strain with respect to the effect of age and CR on the microbiome.

**Table 1:**
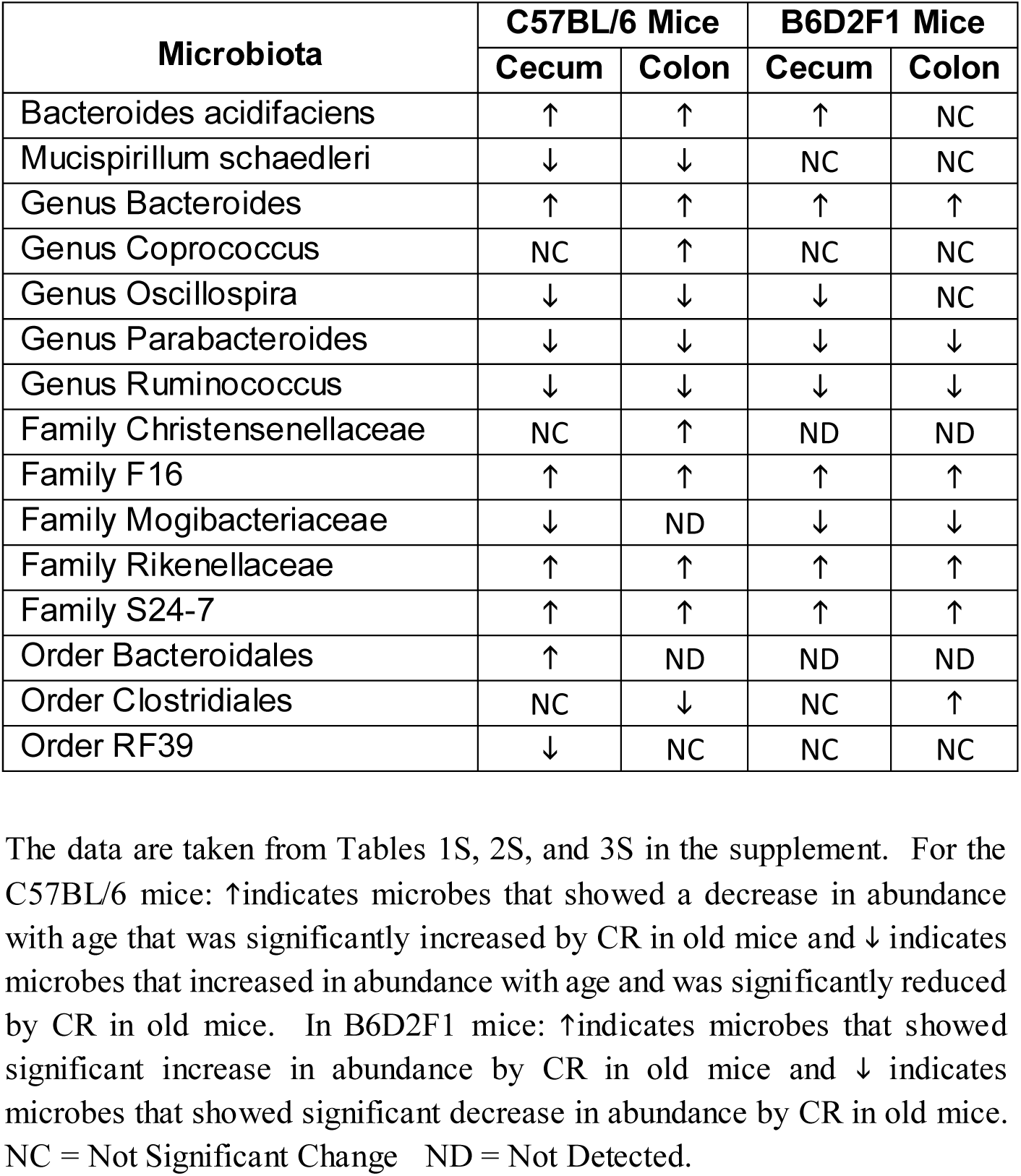
Microbiota that Changed with Age and are Reversed by Caloric Restriction

| Microbiota | C57BL/6 Mice |  | B6D2F1 Mice |  |
| --- | --- | --- | --- | --- |
|  | Cecum | Colon | Cecum | Colon |
| Bacteroides acidifaciens | ↑ | ↑ | ↑ | NC |
| Mucispirillum schaedleri | ↓ | ↓ | NC | NC |
| Genus Bacteroides | ↑ | ↑ | ↑ | ↑ |
| Genus Coprococcus | NC | ↑ | NC | NC |
| Genus Oscillospira | ↓ | ↓ | ↓ | NC |
| Genus Parabacteroides | ↓ | ↓ | ↓ | ↓ |
| Genus Ruminococcus | ↓ | ↓ | ↓ | ↓ |
| Family Christensenellaceae | NC | ↑ | ND | ND |
| Family F16 | ↑ | ↑ | ↑ | ↑ |
| Family Mogibacteriaceae | ↓ | ND | ↓ | ↓ |
| Family Rikenellaceae | ↑ | ↑ | ↑ | ↑ |
| Family S24-7 | ↑ | ↑ | ↑ | ↑ |
| Order Bacteroidales | ↑ | ND | ND | ND |
| Order Clostridiales | NC | ↓ | NC | ↑ |
| Order RF39 | ↓ | NC | NC | NC |

The data are taken from Tables 1S, 2S, and 3S in the supplement. For the C57BL/6 mice: ↑ indicates microbes that showed a decrease in abundance with age that was significantly increased by CR in old mice and ↓ indicates microbes that increased in abundance with age and was significantly reduced by CR in old mice. In B6D2F1 mice: ↑ indicates microbes that showed significant increase in abundance by CR in old mice and ↓ indicates microbes that showed significant decrease in abundance by CR in old mice. NC = Not Significant Change ND = Not Detected.

**Figure 1.**
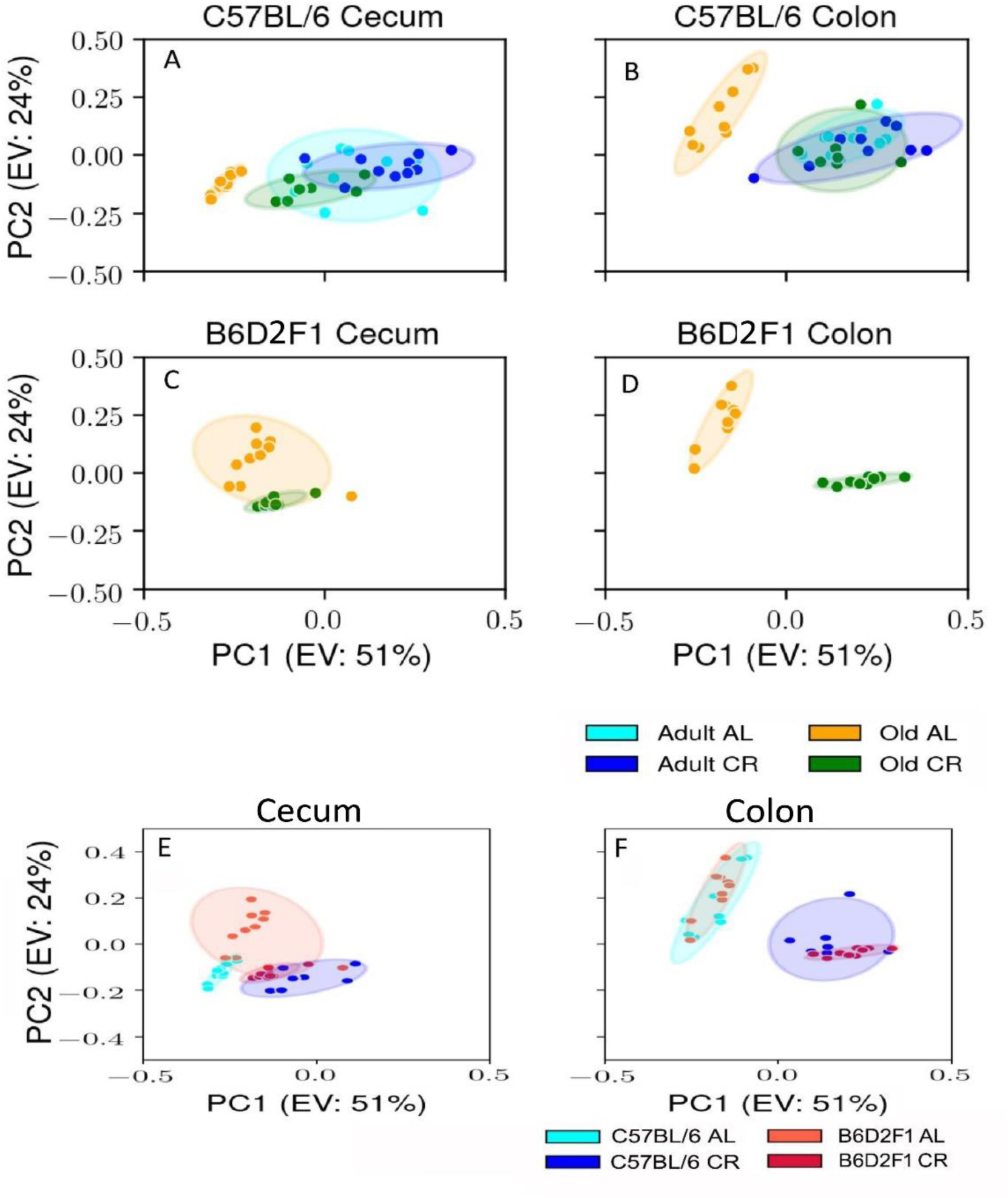
PCA plots showing the variance in the microbiome of mice fed AL and CR. The PCA plots are shown for the microbiome from the cecum (A) and colon (B) of adult and old C57BL/6JN mice fed AL or CR and from the cecum (C) and colon of old B6D2F1 mice fed AL or CR. Panels E and F show PCA plots of the microbiome from the cecum and colon, respectively for old C57BL/6JN and B6D2F1 mice fed AL and CR. Ellipses are 95% confidence intervals for the group obtained using bootstrapping, and “EV: XX%” in the axis labels is the percentage of explained variance of the component.

We next studied the effect of age and CR on the diversity of the microbial species present in the microbiome because microbiota diversity is often used as a measure of the ‘health’ of the microbiome, the higher the ratio the ‘healthier’ the microbiome. The data in Figure 2A and B show the diversity of the microbiota from cecum and colon of C57BL/6JN mice using Shannon’s diversity index, which equally weights richness (number of different taxa) and evenness (equitability of taxa frequencies) of the microbiome (Wagner et al., 2018). The diversity of the microbiome in the cecum and colon decreased with age in C57BL/6JN mice fed AL; however, the decrease was only significant for the colon. CR had no effect on the diversity of the microbiome in adult mice; however, in old mice, CR resulted in a significant increase in the diversity of the microbiome in both the cecum and colon, which was comparable to the diversity observed in the adult mice. Figure 2C shows the diversity of the microbiota of the cecum and colon from the old B6D2F1 mice fed AL or CR. In B6D2F1 mice, CR had no significant effect on the diversity of the microbiota in the cecum; however, there was a significant increase in the diversity of the microbiota of the colon.

**Figure 2.**
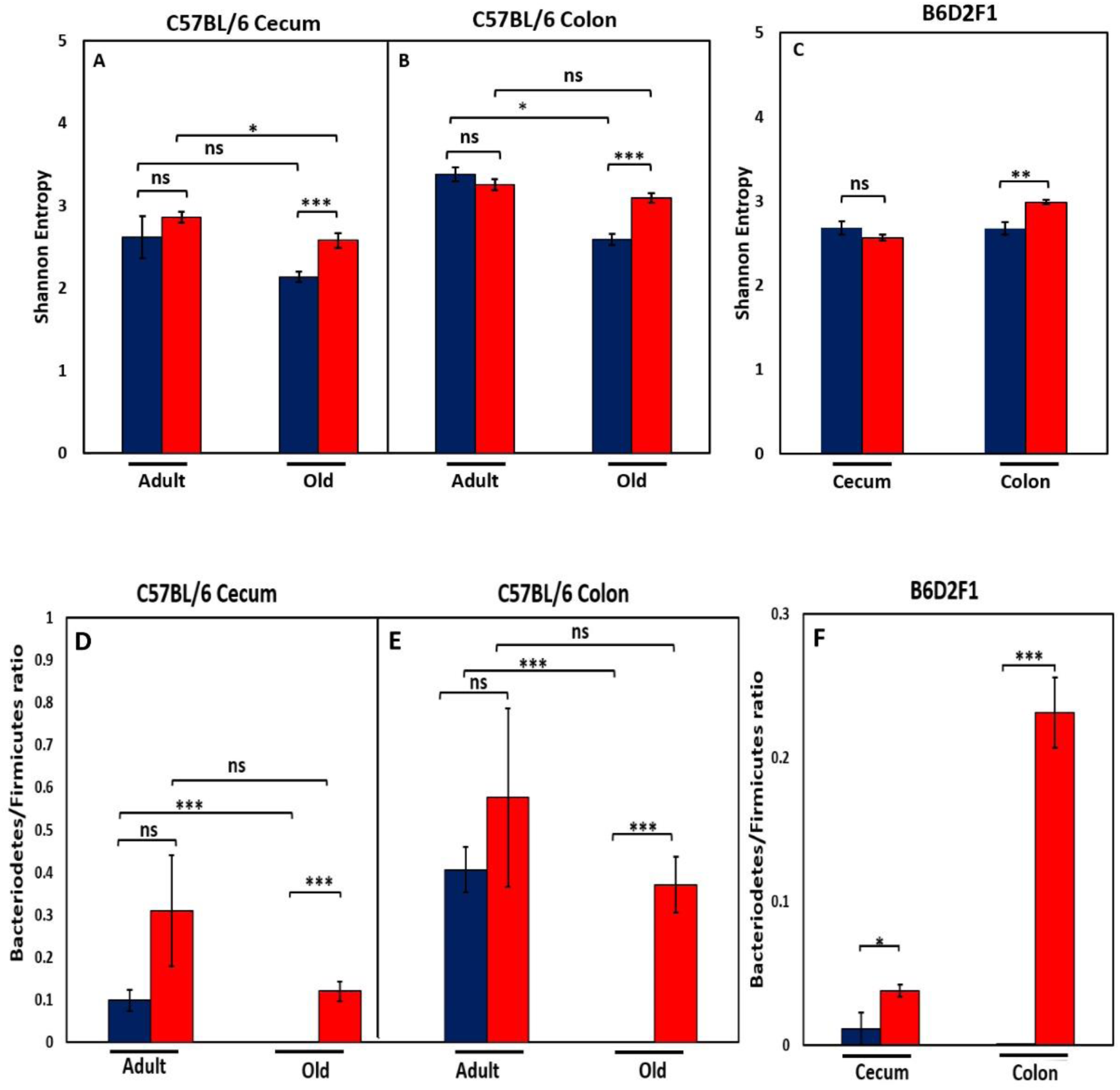
Effect of age and CR on microbiome. The top panel show the Shannon diversity index (Wagner et al., 2018) for the cecum (A) and colon (B) of adult and old C57BL/6JN mice and old B6D2F1 mice (C) fed AL (blue bars) or CR (red bars). The bottom panel show the ratio of *Bacteriodetes* to *Firmicutes* for the cecum (D) and colon (E) of adult and old C57BL/6JN mice and for the cecum and colon of old B6D2F1 (F) mice fed AL (blue bars) or CR (red bars). The data represent the mean and SEM for 8-10 mice per group. *p<0.05, **p<01, ***p<0.001, ns = non-significant.

We also measured the ratio of *Bacteriodetes* to *Firmicutes* in the cecum and colon of the C57BL/6JN and B6D2F1 mice because this ratio has been correlated with various diseases (Verdam et al., 2013; Ley et al., 2006) and has been reported to change with age (Mariat et al., 2009; Lee et al., 2018). Figure 2D and E show that the ratio of *Bacteriodetes* to *Firmicutes* decreases in both the cecum and colon with age in C57BL/6JN mice, and this decrease is attenuated by CR in old mice resulting in a ratio similar to adult mice fed AL. Although CR increased the ratio of *Bacteriodetes* to *Firmicutes* in adult mice, this increase was not statistically significant. In the B6D2F1 mice (Figure 2F), the ratio of *Bacteriodetes* to *Firmicutes* was significantly higher in CR mice compared to mice fed AL especially in the colon.

The relative abundance of all the microbes found in the cecum and colon of C57BL/6JN and B6D2F1 is presented in Figures 1S and 2S in the supplement. Figures 3 and 4 show the abundance of microbes that changed significantly with age or diet for the C57BL/6JN and B6D2F1 mice, respectively. From Figure 3, it is obvious that a major age-related change occurs in the microbiota of C57BL/6JN mice at all three taxonomical levels, and this difference is apparent in both the cecum and colon. The data in Figure 3 also show that CR had only a small effect on the microbiota in the cecum and colon of adult mice. However, CR in the old mice showed a dramatic change in the microbiota compared to old AL in both the cecum and colon, with the microbiota becoming more like the adult C57BL/6JN mice, which is consistent with the PCA data in Figure 1. At the family level, the adult AL and CR mice and old CR mice had similar microbial patterns with a greater increased abundance in the S24-7 and Rikenellaceae families. In contrast, old AL mice showed a higher abundance of the Lachnospiraceae and Ruminococcaceae families. At the genus level, the microbial pattern was quite different for the cecum and colon in contrast to the family and species levels. In the cecum, Bacteroides was the prominent genus in the adult AL and CR mice and old CR mice while Parabacteroides and Lactobacillus predominated in the old AL mice. In the colon, Oscillospira was the prominent genus in all four groups of mice. At the species level, Bacteroides acidifaciens was the most abundant microbe (>60%) in adult AL and CR mice and old CR mice, whereas Ruminococcus gnavus and Muscispirillum schaedleri were more abundant in the old AL mice in both the cecum and colon.

**Figure 3:**
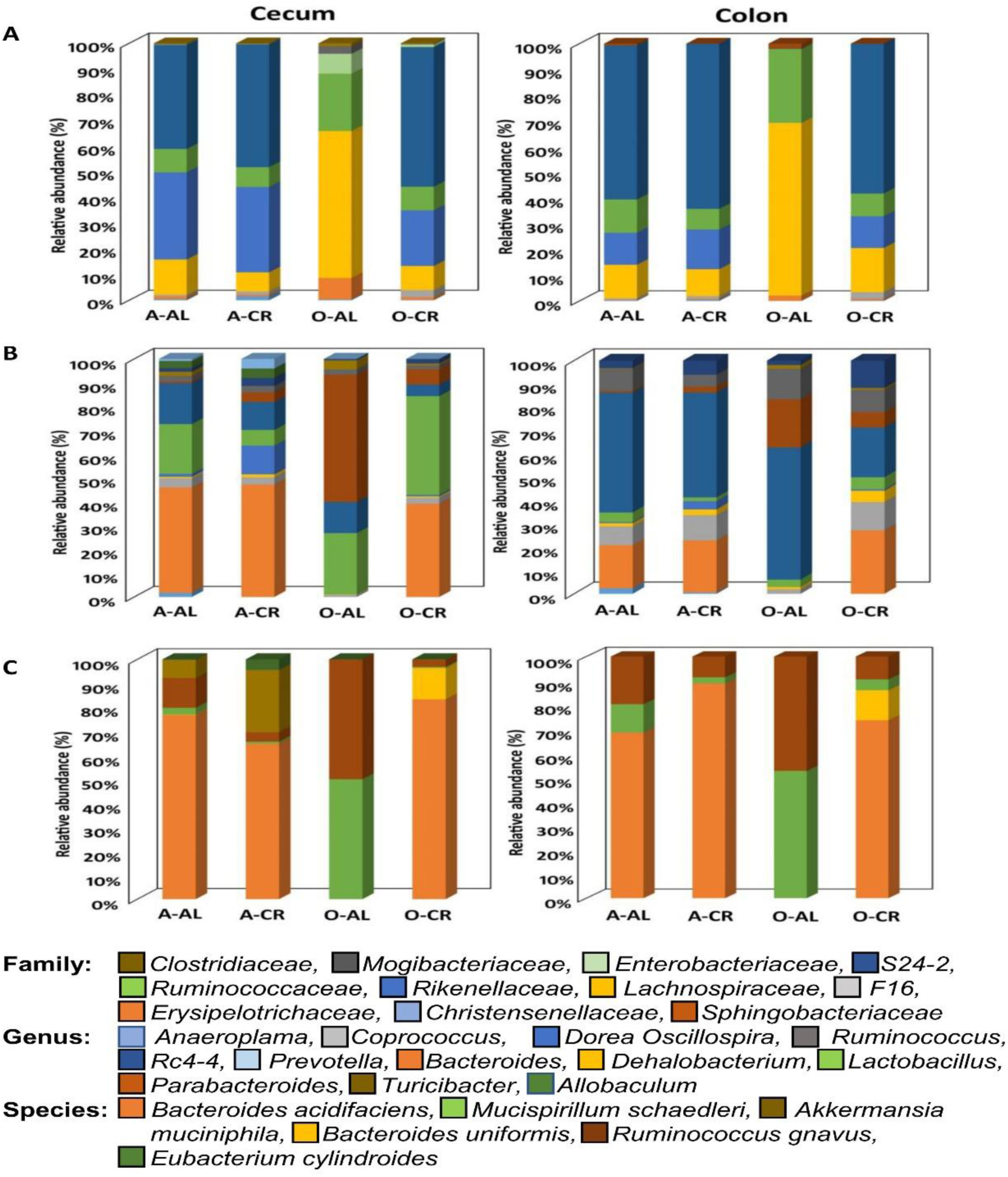
The abundance of microbes that changed with age or CR in C57BL/6 mice. The relative abundance of the 34 microbes that changed significantly in C57BL/6JN mice are shown for adult mice fed AL (A-AL) or CR (A-CR) and old mice fed AL (O-AL) or CR (O-CR). The microbiome data are presented on the basis of family (A), genus (B), or species (C) of the microbes found in either the cecum or colon of the mice.

**Figure 4:**
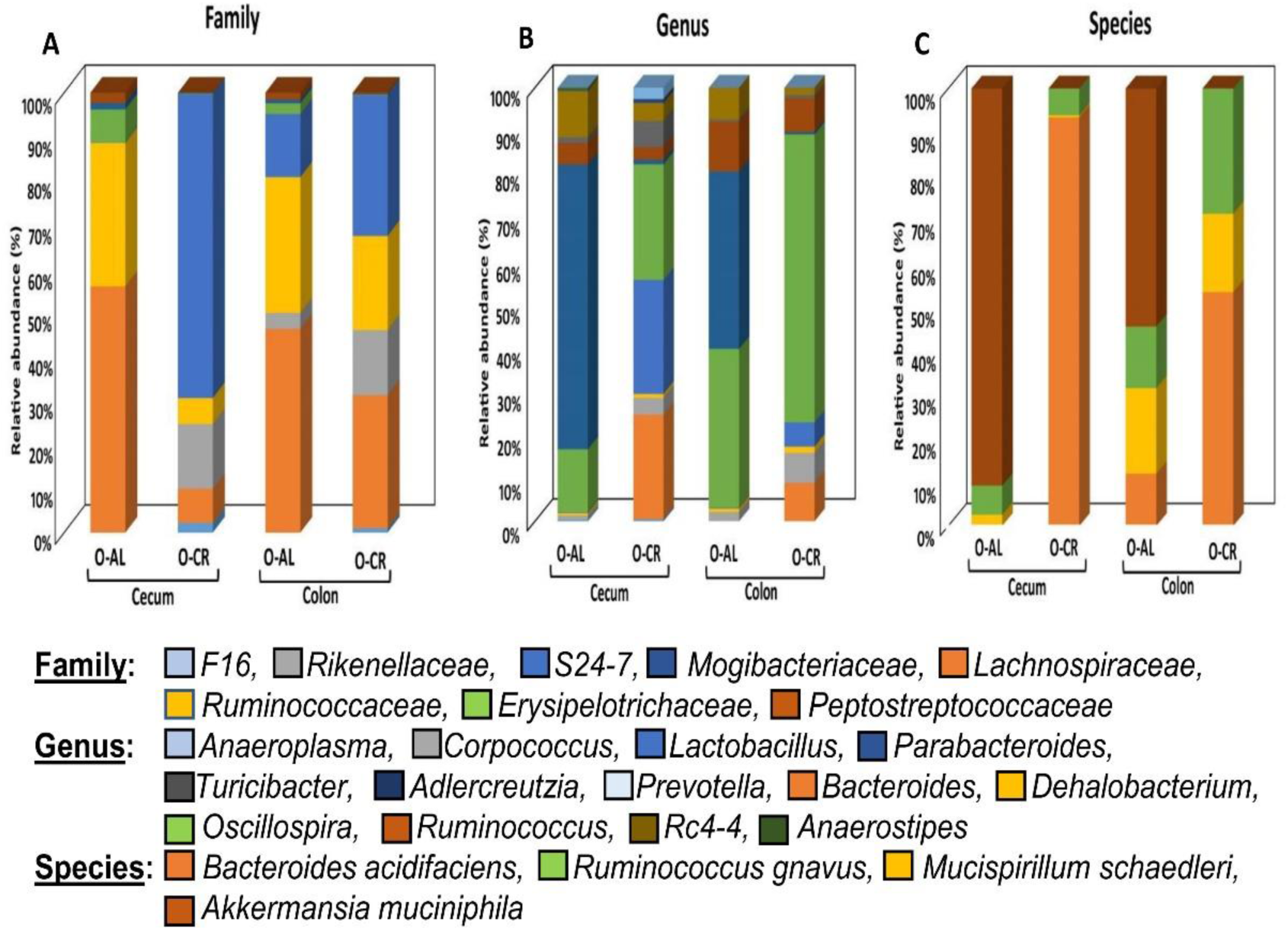
The abundance of microbes that changed with CR in B6D2F1 mice. The relative abundance of the 27 microbes that changed significantly in B6D2F1 mice are shown for old mice fed AL (O-AL) or CR (O-CR). The microbiome data are presented on the basis of family (A), genus (B), or species (C) of the microbes found in either the cecum or colon of the mice.

Figure 4 shows the relative abundance of the microbes that changed with age or diet for old B6D2F1 mice. At the family level, the cecum and colon from old AL showed similar profiles with a greater abundance of Lachnospiraceae and Ruminococcaceae as was found in C57BL/6JN mice. The cecum of old CR B6D2F1 mice showed a greater abundance of S24-7 and Rickenellaceae families (which was similar to C57BL/6JN mice) whereas the colon of old CR B6D2F1 mice showed an abundance in the S24-7, Lachnospiraceae, Ruminococcaceae and Rickenellaceae families. At the genus level, the cecum and colon of the old AL B6D2F1 mice showed an increase in the abundance of the Parabacteroides, Oscillospira, Ruminococcus and Rc4-4 families. The cecum of old CR mice showed a greater abundance in Lactobacillus, Oscillospira and Bacteroides families whereas the colon of old CR mice had increased abundance of Oscillospira (>50%). In the Cecum and colon of old B6D2F1 mice fed AL, Akkermansia municiphila was the prominent species found. In contrast, Bacteroides acidifaciens and Ruminococcus gnavus were the predominant species in the cecum and colon of old CR mice.

Of the 35 microbes from different taxonomic levels identified in the gut of C57BL/6JN mice (Tables 1S and 2S), 15 showed a significant change with age that was attenuated by CR in old animals, and these 15 are listed in Table 1. The number (8) of microbes that increased with age and were reduced by CR is similar to the number (7) of microbes that showed a decrease with age and were increased by CR. It is interesting to note, that CR significantly altered the abundance of 11 of the 35 microbes in the cecum of adult mice; 5 of the microbes were significantly altered by CR in old mice. In the colon, the abundance of only one microbe (Christenellaceae family) was significantly altered by CR in adult mice. Thirteen of the 15 microbes listed in Table 1 were also detected in B6D2F1 mice, and of these 13 microbes, over 75% (10) showed the same changes in abundance with CR as was observed in the C57BL/6JN mice. However, CR significantly altered the abundance of several microbes in the old B6D2F1 mice that were not observed in old C57BL/6JN mice. For example, CR reduced the abundance of Akkermansia muciniphila, the rd4-4 genus, and the Erysipelotrichaceae family in the colon of the B6D2F1 mice. In the cecum of the B6D2F1 mice, CR altered the abundance of 17 of the 27 microbes identified. For both C57BL/6JN and B6D2F1 mice, CR altered significantly the abundance of more microbes in the cecum than in the colon.

Figure 5 shows the effect of age and CR in C57BL/6JN mice on the abundance of the six microbes from Table 1 that were the most abundant; collectively, these microbes comprised between 5 to 50% of the microbiome. The abundance of all six microbes showed similar changes with CR in the cecum and colon. Four of microbes showed a major decrease in abundance with age (Bacteroides acidfaciens, Bacteroides, Rikenellaceae, and S24-7), which was completely reversed by CR. Two of the most abundant microbes that showed the most dramatic changes were the genus Parabacteroides and the S24-7 family. Parabacteroides increased 100-fold in the cecum from less than 1% to over 25% of the microbiota in the old AL mice. In the colon, Parabacteroides abundance increased over 30-fold in the colon making up almost 5% of the microbiota in the old AL mice. CR reduced the abundance of Parabacteroides to less than 2% in both the cecum and colon of old mice. The S24-7 family of microbes decreased from 17 to 27% in the cecum and colon, respectively, of adult mice to negligible levels (less than 0.2%) in the old AL mice. CR resulted in an increase in the abundance of the S24-7 family to over 19% in the cecum and colon.

**Figure 5.**
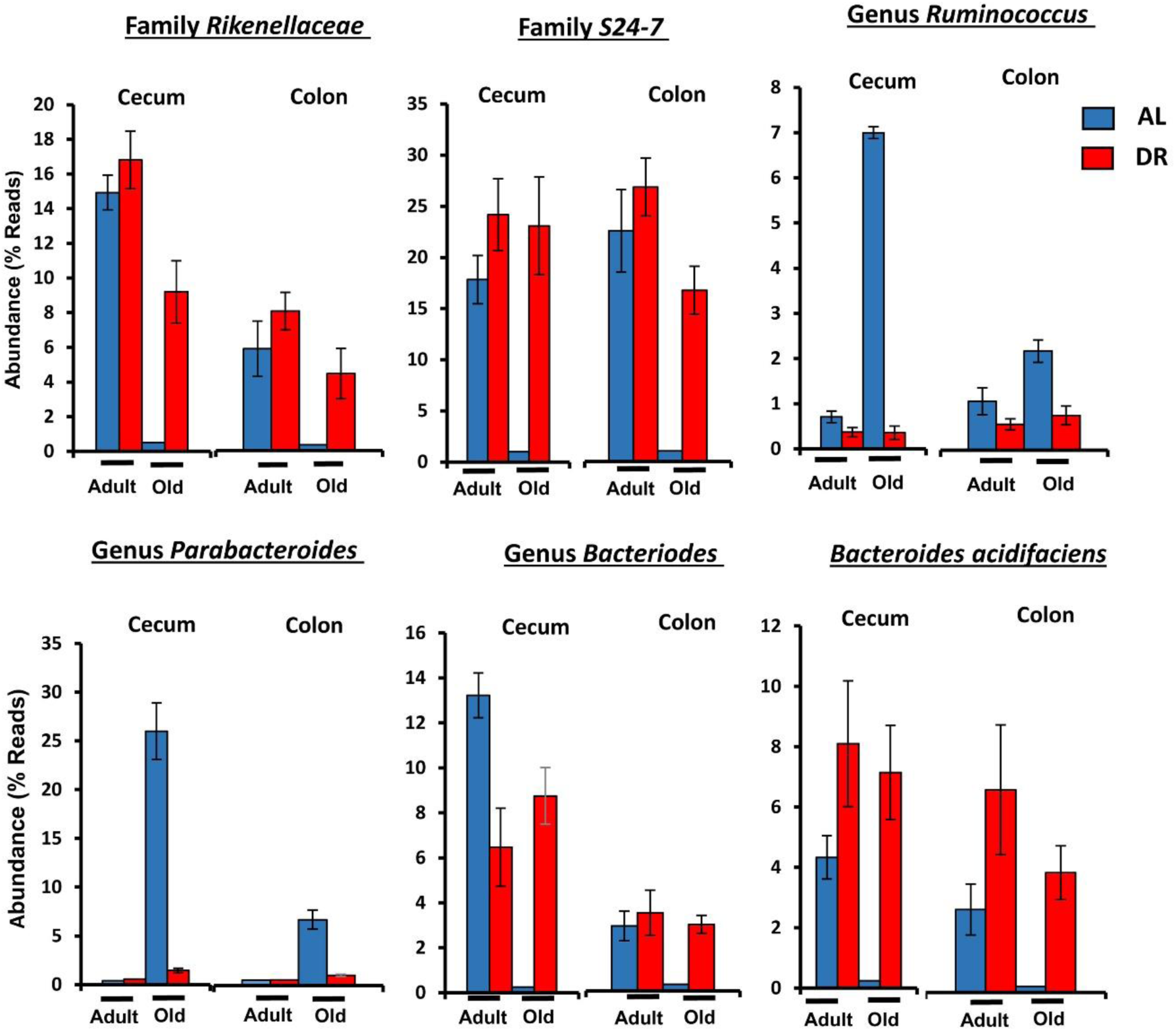
The effect of age and CR on the abundance of the six major microbes found in C57BL/6JN mice. The relative abundance (mean and SEM) of six microbes from Table 1 are shown for 8-10 mice per group for adult and old C57BL/6JN mice fed AL (blue bars) and CR (red bars).

We next determined the degree of association in the abundance of the four microbial classes that made up the majority of microbes that changed with age and diet. Figure 6 shows the t-statistic from the OLS model when each coefficient is plotted for each microbial class. A positive t-statistic for diet indicates that microbial abundance in that class increases with CR, and a positive t-statistic for age indicates that microbial abundance increases with age. The data in Figure 6 show a negative correlation for the phylogenetic class with CR and age, indicating that CR has opposing effects on the gut microbiome compared to age. In addition, the phylogenetic class Bacteroidia is associated with CR and youth, whereas the class Clostridia is associated with AL and age.

**Figure 6.**
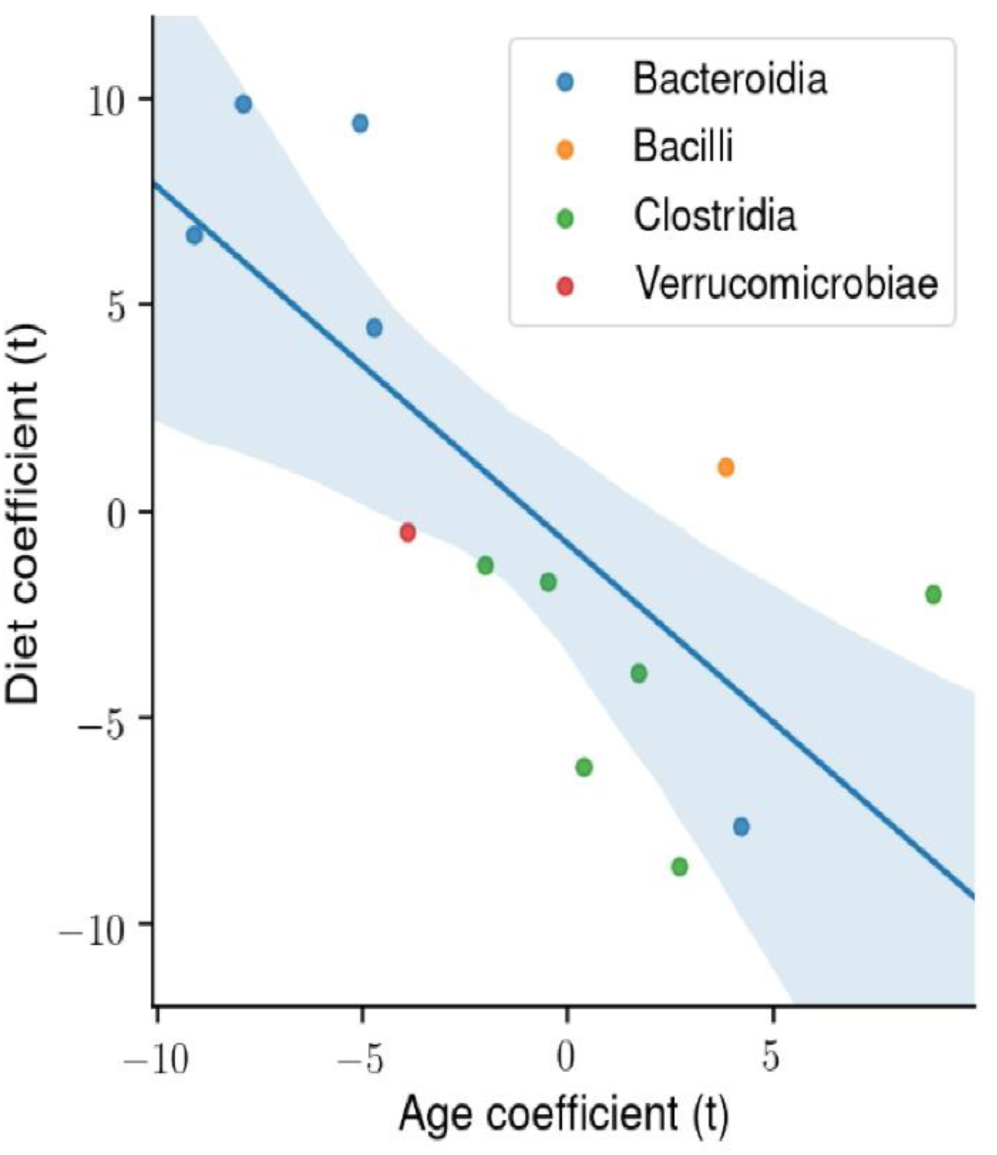
The degree of association between microbial taxa abundance, age, and diet. Using ordinary least squares (OLS) linear model, the t-statistic for each coefficient is plotted for the microbial phylogenetic classes: *Bacteroidia, Bacilli, Clostridia, Verrucomicrobiae*. There is a significant negative correlation between the age and diet t-statistic.

### Effect of age and caloric restriction on the metabolite composition of the fecal material from the colon of mice

To understand how age and CR might affect the microbiome, we also measured the metabolites in the fecal pellets from the colon of the C57BL/6JN and B6D2F1 mice by targeted liquid chromatography mass spectrometry as described in the Methods. Table 4S in the supplements lists the 98 metabolites detected in the fecal samples and the levels of these metabolites in the fecal pellets obtained from the two strains of mice. We observed that the levels of 57 of the 98 metabolites exhibited a significant change with either age (old AL vs adult AL in C57BL/6JN) or CR in the C57BL/6JN (old AL vs old CR) and B6D2F1 (old AL vs old CR) mice as shown in the Venn diagram in Figure 7A. The levels of 36 metabolites changed with age in AL C57BL/6JN mice, 30 changed with CR in the old C57BL/6JN mice, and 43 changed with CR in the old B5D2F1 mice. Approximately 60% of the metabolites that changed with age were significantly attenuated by CR in the C57BL/6JN mice and two-thirds of the metabolites that changed with CR were the same for the old C57BL/6JN and B6D2F1 mice. The data in Table 4S were analyzed by principal component analysis, and the results are shown in Figure 7B and C. For the C57BL/6JN mice, the adult AL and CR mice and the old CR mice overlapped; however, the old mice fed AL formed a separate cluster (Figure 7B). Figure 7C shows the PCA for old mice fed either AL or CR. While there was some overlap between old C57BL/6JN and B6D2F1 mice fed AL, the old CR C57BL/6JN or B6D2F1 mice were clearly separated from the old mice fed AL, with the B6D2F1 mice showing the greatest separation.

**Figure 7:**
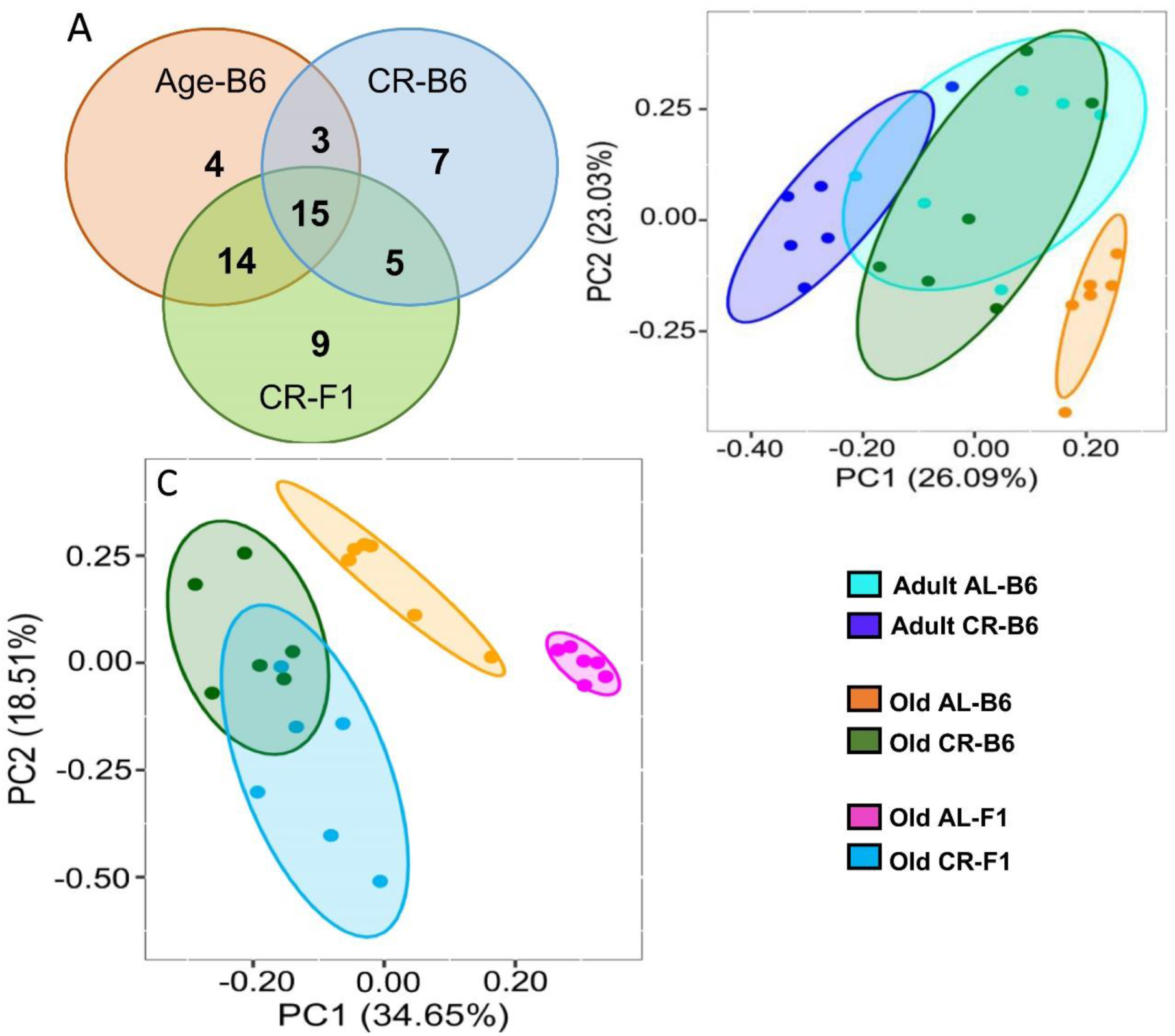
Effect of age and CR on the fecal metabolome. **Panel A:** Venn diagram showing the distribution of the number of metabolites that changed significantly in C57BL/6JN mice with age (orange) or with CR in old C57BL/6JN mice (blue) or changed significantly with CR in B6D2F1 mice (green). **Panel B:** PCA Analysis of fecal metabolites from adult and old C57BL/6JN mice fed AL or CR (6 mice/group). Data normalized and imputed from the 98 metabolites. **Panel C:** PCA Analysis of fecal metabolites from old AL and CR C57BL/6JN and B6D2F1 mice (6 mice/group) normalized and imputed from the 98 metabolites.

Figure 8A shows the heatmap of the 48 metabolites that changed significantly (FDR <0.05) in the C57BL/6JN mice (Old AL vs Adult AL and Old AL vs Old CR). It is evident, that the pattern of the fecal metabolites from the old AL mice was quite different from that observed for the adult AL or CR mice. The pattern of metabolites in the old CR approached a pattern more similar to that observed in the adult mice. Figure 8B shows the heatmap of the 53 metabolites that were changed significantly by CR (Old AL vs Old CR) in either old C57BL/6JN mice or old B6D2F1 mice. The pattern of fecal metabolites is quite different for the old C57BL/6JN and B6D2F1 mice fed even though their microbiome composition were similar (Figure 1F). CR resulted in a dramatic change in the pattern of metabolites in both strains of mice. Interestingly, the difference in the metabolite pattern between the C57BL/6JN and B6D2F1 mice was largely resolved by CR. CR had a greater effect on the fecal metabolome of the B6D2F1 mice than the C57BL/6JN mice.

**Figure 8:**
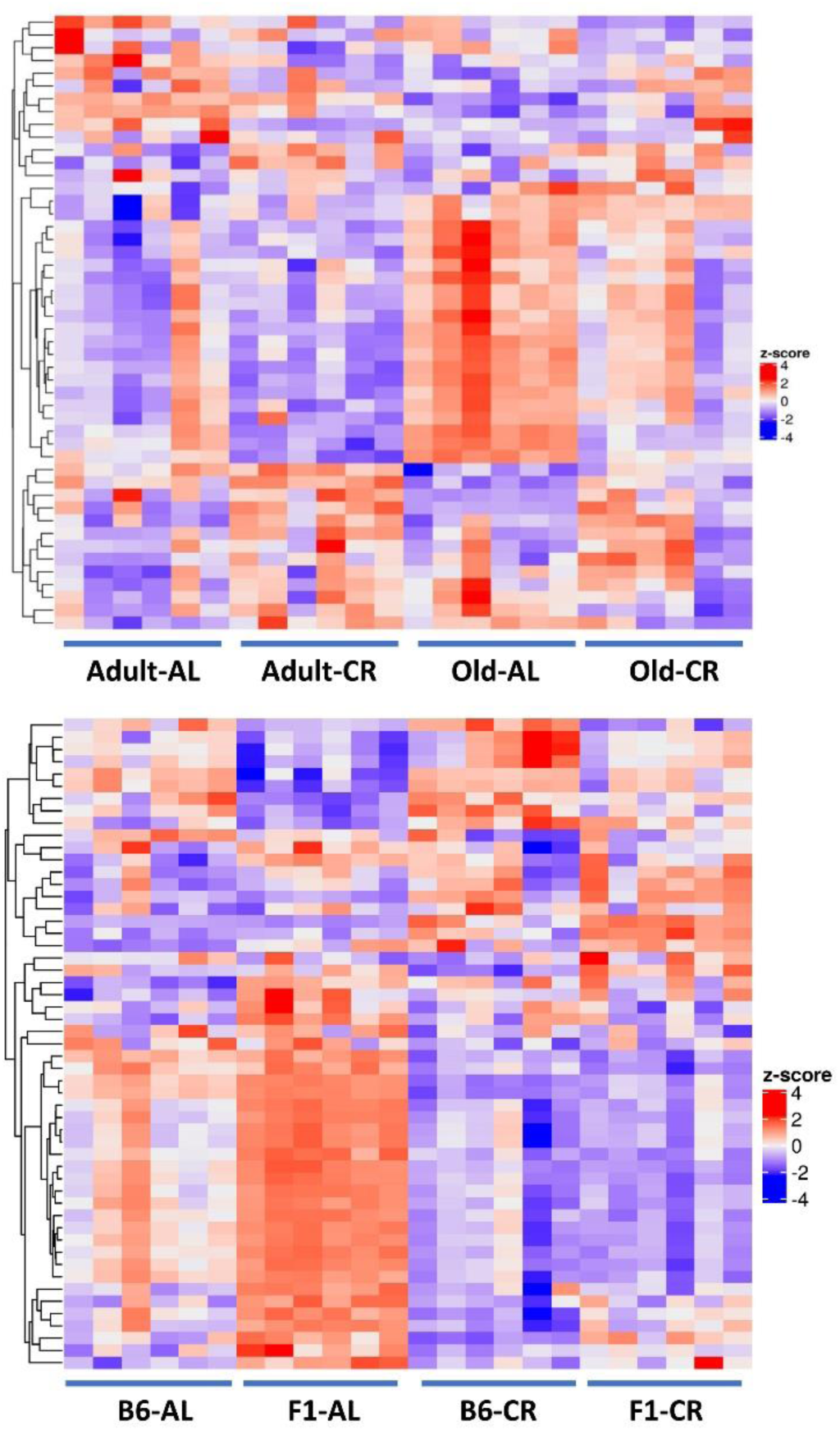
Heatmaps of the fecal metabolome. **Top Panel:** Heatmap of the 48 metabolites that changed significantly (FDR <0.05) with either Old AL vs. Adult AL or Old AL vs. Old CR comparisons in C57BL/6JN mice (6 mice per group). **Bottom Panel:** The heatmap shows the relative levels of the 53 metabolites that changed significantly (FDR <0.05) with CR in either old C57BL/6JN or old B6D2F1 mice (6 mice per group). In order to generate the heatmap, we computed a z-score of the log2-abundance, where we adjusted the data, by metabolites, to have a mean of zero and a standard deviation of 1. The heatmap is generated using the ComplexHeatmap R package, where the metabolites were clustered via the hclust function with the “complete” agglomeration method. Distance matrix for clustering are computed using “Euclidean” distance. The resulting heatmap presents the metabolites in rows and samples in columns.

We next determined the effects of age, diet, and strain on the metabolomic profile using an OLS linear model. The coefficients of this model reflect the independent fold changes on metabolic pathways represented by the metabolites that changed with age, diet, and strain. The heatmap in Figure 9A shows the average log2FC attributable to age, diet, and strain among all the metabolites measured in the respective pathways. The data show the following: the metabolomic effects of age and DR are broadly anti-correlated and the metabolomic profile of the B6D2F1 mice is correlated with that of CR and anti-correlated with that of age, suggesting that the B6D2F1 mice have a ‘younger’ and more ‘CR-like’ metabolomic profile compared to the C57BL/6JN mice. To contrast the effects of age and CR on the metabolomic profile, we again used an OLS linear model where the coefficients reflect the independent fold changes in metabolome abundance resulting from age and diet. Figure 9B shows the average log2FC attributable to age and diet among all the metabolites measured in their respective pathways. The data show that the effects of age and CR on the metabolome are broadly anti-correlated as one would predict. In addition, pathways related to amino acid metabolism broadly increase with age and decrease with CR, whereas pathways associated with nucleotide metabolism increase with CR and decrease with age.

**Figure 9:**
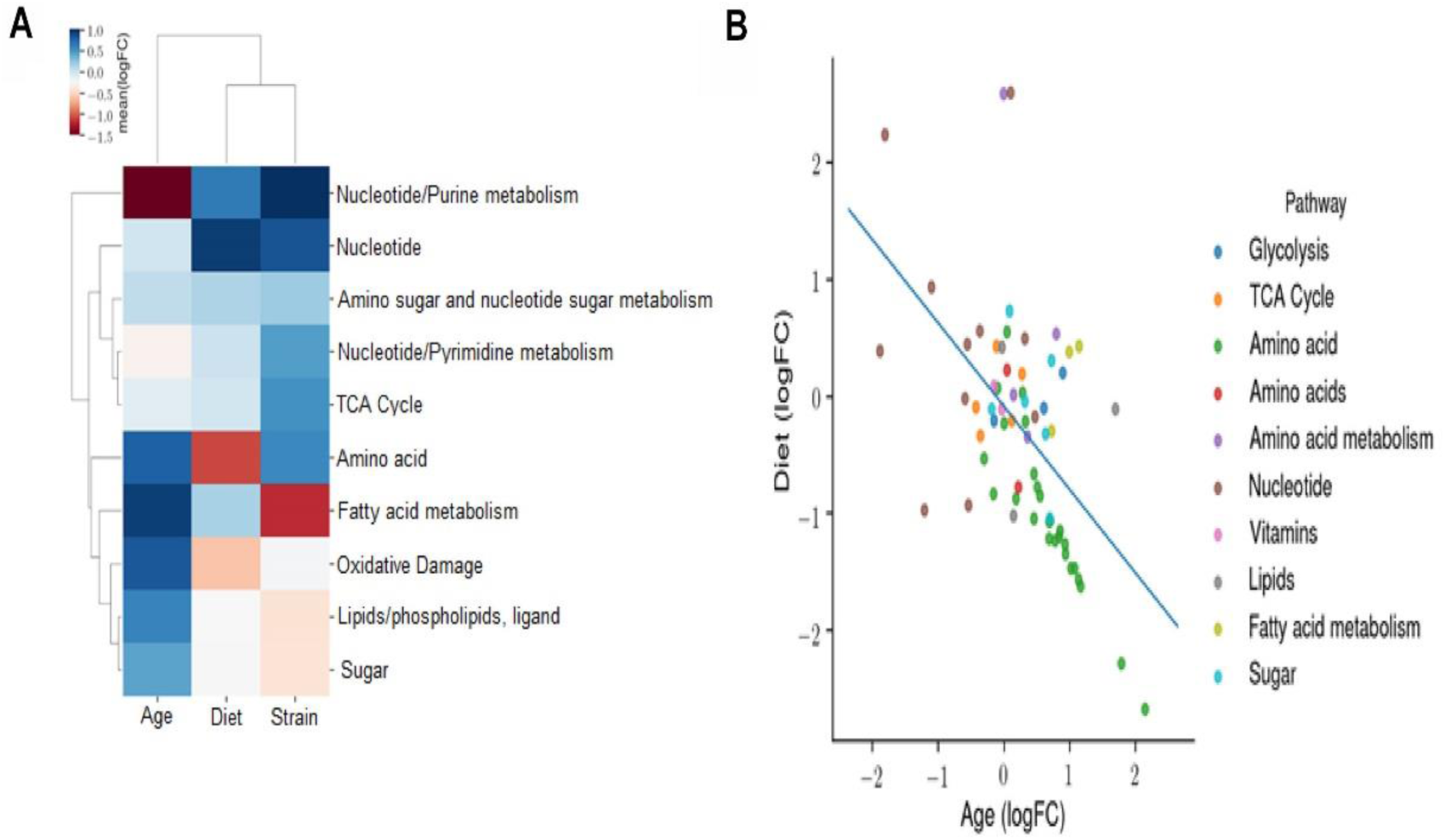
Effect of age, diet, and strain on the fecal metabolome. **Panel A:** An ordinary least squares (OLS) linear model with the formula *log(Abundance) ∼ Age + Diet + Strain* is fitted to evaluate the effects of age, diet, and strain on the metabolomic profile. The coefficients of this model reflect the independent fold changes on metabolomic abundances caused by age, diet, and strain. This panel displays the average log_2_FC attributable to age, diet, and strain among all the metabolites measured in the respective pathways. A positive value for age, diet, or strain indicates that this metabolomic pathway is increased with age, DR, or B6D2F1 (versus C57BL/6JN), respectively. **Panel B:** OLS linear model coefficients reflecting the independent fold changes in metabolomic abundances caused by age and diet. The average log_2_FC attributable to age and diet among all the metabolites measured in the respective pathways are shown. A positive value for age or diet indicates that this metabolomic pathway is increased with age or CR, respectively.

### Effect of age and caloric restriction on the transcriptome of mucosa from the colon of mice

To gain an understanding of the potential interaction between the microbiome and intestine, we studied the effect of age and CR on the transcriptome of intestinal mucosa isolated from the colon of C57BL/6JN mice used to study the microbiome and metabolome. We identified a total of 45,796 transcripts, and after filtering for low raw counts, there were 36,083 transcripts. Differential expression analysis showed that the expression of 723 mucosal genes changed significantly [P (Corr) < 0.05 and 2.0-fold] with either age (old AL vs adult AL) or CR (old CR vs old AL and adult CR vs adult AL) in C57BL/6JN mice. Of the 660 genes that changed with age (279 increased, 381 decreased), CR reversed the age-related changes in 189 of the genes (29%) in old mice. In addition, CR significantly changed the levels of 57 genes that did not change with age in old mice, i.e., approximately one-fourth of the genes that changed with CR did not change with age. Interestingly, CR resulted in significant changes in 94 of the genes in adult mice and over 50% of these genes remained changed by CR in the old mice. Table 5S in the supplements lists the top 20 genes whose expression changed the most with age and CR.

Figure 10A shows PCA analysis of differentially expressed genes [P (Corr) < 0.05 and 2.0-fold] in old CR, old AL, adult CR based on adult AL in the C57BL/6JN mice. The transcriptomes of the adult and old CR mice formed clusters that were clearly separated from each other. In contrast to what we observed for the microbiome and metabolome, both adult CR and old CR formed clusters that were clearly separate from their AL counterparts. Interestingly, adult and old CR mice formed clusters that overlapped, suggesting a similarity in the transcriptome of adult and old CR mice. Figure 10B shows the heatmap of genes that changed significantly in the C57BL/6JN mice. CR resulted in a major shift in the transcriptome pattern in both adult and old mice. These data agree with the PCA data in Figure 10A, which showed that CR altered the transcriptome in both adult mice and old mice.

**Figure 10:**
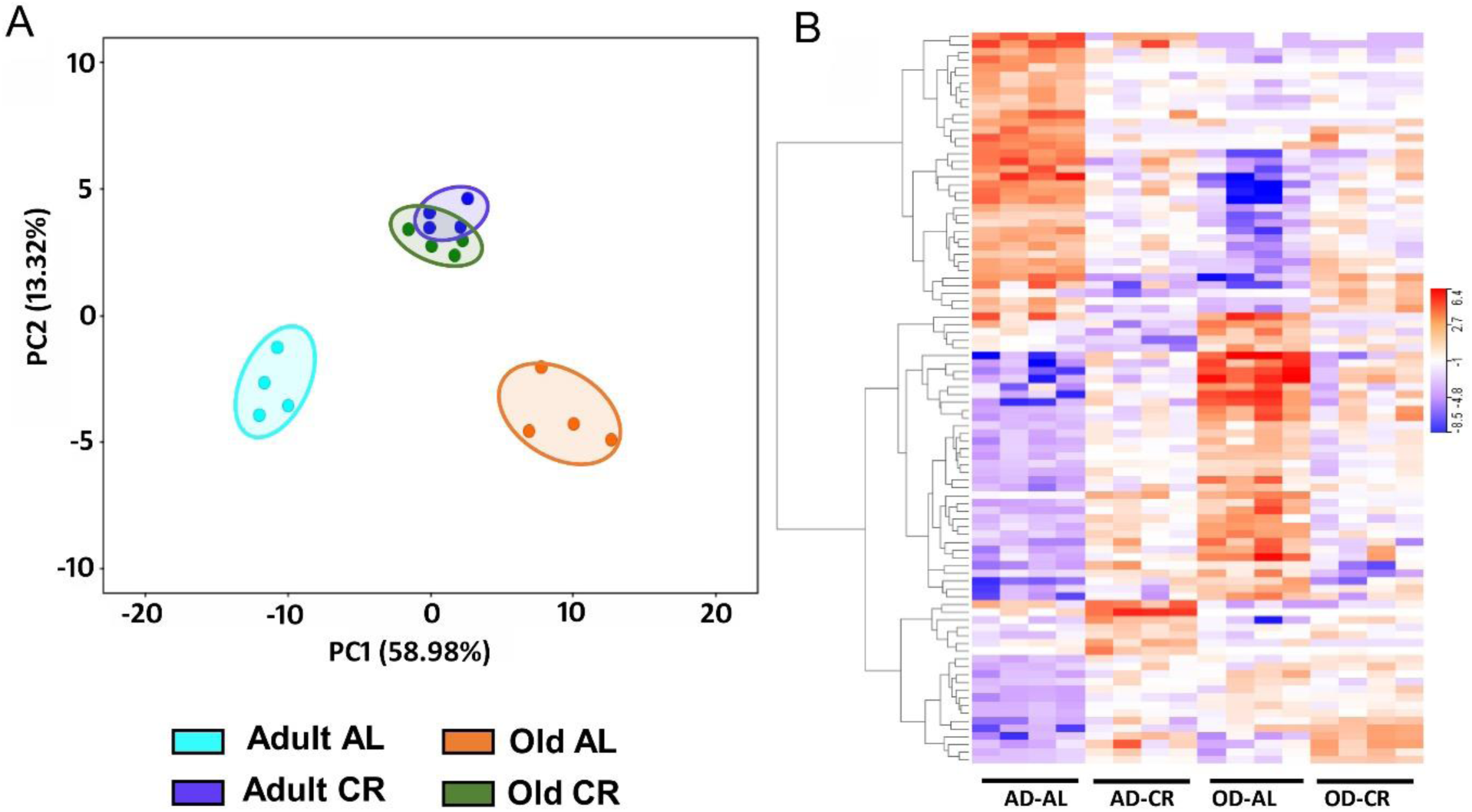
Analysis of colon mucosa transcriptome of C57BL/6JN mice fed AL and CR. The data were collected from 4 mice per group, and the PCA and heatmap plots show those transcripts that show 2-fold change in expression [P (Corr) <=0.05). **Panel A:** PCA analysis of transcriptome from adult and old mice fed AL or CR. **Panel B:** Heatmap of the relative levels of the transcripts that changed significantly with either age and/or CR in individual mice compared to Adult AL mice.

## DISCUSSION

The first study to show that life-long CR (30% started at 5 weeks of age) significantly changed the gut microbiota of old (15 and 35 months) mice was reported by Zhang et al (2013). Subsequently, Kok et al (2018) also reported that life-long CR (30% started at 9 weeks of age) significantly changed the microbiome of old (28 months) mice. However, in a study focusing primarily on response to influenza, Bartley et al (2017) reported that CR (40% starting at 3-4 months of age) had very little effect on the microbiome of old (19 to 21 months) mice. These three studies used male C57BL/6 mice. The studies by Zhang et al (2013) and Kok et al. (2018) were conducted on C57BL/6 obtained and maintained in either China or The Netherlands, respectively, while Bartley et al (2017) used C57BL/6JN mice obtained from the animal colony maintained by NIA. Because environment can have a major impact on the gut microbiome (Ericsson et al., 2018) and the NIA animal colony is maintained under optimal husbandry and used by investigators across the country, we believed it was important to characterize in detail the effect of CR on the microbiome in mice from the NIA animal colony microbiome. In this study, we evaluated the effect of 40% CR started at 3-4 months of age on the microbiota of both the cecum and colon of 9- and 24-month-old C57BL/6JN mice, i.e., mice were fed a CR-diet for either 5 or 20 months. This is the first study to assess the effect of age or CR on the cecum and to simultaneously compare the effect of short term and life-long CR on the microbiome. Our data show that both age and CR brought about significant changes in the microbiome in both the cecum and colon of C57BL/6JN mice. Most interestingly, CR attenuated the age-related changes in microbiome and resulted in a microbial profile of old mice becoming similar to that of adult mice. For example, CR altered the microbial diversity in both cecum and colon, which has been associated with optimal health and has been shown to change with age (Cuesta-Zuluaga et al., 2019). In addition, healthy, elderly have been shown to have a diverse gut microbiome compared to elderly individuals with various comorbidities, e.g., dysbiosis and reduced microbial diversity (Deng et al., 2019). We found that CR increased diversity in the cecum and colon of old mice, which could potentially help maintain gut integrity and homeostasis. When we analyzed the microbial flora, the two most prominent phyla identified were the gram positive Firmicutes and gram negative Bacteriodetes. Studies in animals (Ley et al., 2005) and humans (Koliada et al., 2017) have shown that reduced levels of Bacteriodetes and increased Firmicutes are associated with obesity. Our data show that CR increased the Bacteriodetes/Firmicutes ratio in both the cecum and colon C57BL/6JN mice and this observation tracks with the fact that CR reduces adiposity and improves lean body mass (Matyi et al., 2018).

Comparisons of relative abundance of microbes also showed that age and CR significantly shifted the microbial profile of the intestine. The prominent microbial family observed in the cecum and colon of the old mice fed AL were Lachnospiraceae and Ruminococcaceae, which belong to the phylum Firmicutes. Many microbes in the Lachnospiraceae family are associated with impaired glucose metabolism, obesity, inflammation, and metabolic syndrome (Kameyama and Itoh, 2014; Ai et al., 2019; Vacca et al., 2020). In contrast, the Ruminococceae family includes microbes that are positively correlated to oxidative stress, inflammation, inflammatory bowel diseases, and diabetes (Hall et al., 2017; Henke et al., 2019; Rinninella et al., 2019). Apart from the Lachnospiraceae and Ruminococceae families, we also found the Erysipelotrichaceae and Enterobacteriaceae families, which are reported to be associated with inflammation and infections in the host, to be abundant in the cecum of the old AL mice (Kaakoush, 2015; Bublitz et al., 2014). The abundance of these microbes in old AL mice could play a role in compromising the integrity of the gut. Interestingly, CR altered the relative abundance of several microbes known to have positive health benefits. For example, CR increased the abundance of the Rikenellaceae family and S24-7 family, which belong to the phylum Bacteriodetes. Studies show that the Rikenellaceae family contains hydrogen-producing bacteria that can neutralize reactive oxygen species (Xie et al., 2019) and is inversely associated with inflammation (Kim et al., 2019). In addition, the S24-7 family has been shown to be a fermentative bacteria capable of producing enzymes that breakdown carbohydrates and have positive effects against diabetes (Krych et al., 2015) and inflammatory arthritis (Rogier et al., 2017). The biggest challenge of microbiome studies is that it is hard to conclude whether a shift in microbiota is harmful or beneficial because each microbial family includes bacteria of which some are harmless symbionts and some are harmful pathogens. To add to the complexity, a particular species of bacteria can act both as a ‘good or bad’ bacteria depending on the environment or health of the host (Cirstea et al., 2018). For example, studies have shown that reduction in microbial families such as Lachnospiraceae and Ruminococcaceae are associated with colorectal cancer and inflammatory bowel disorders (Kim et al., 2016). However, our study showed that CR, well known for its protective effect on colon health and colorectal cancer, reduced the abundance of the Lachnospiraceae and Ruminococcaeae families, a scenario that has been reported to be harmful for the gut (Kim et al., 2016).

An important observation in our study was that CR attenuated or reversed the changes that occured with age and changed the intestinal milieu of the old mice to look more like adult mice even down to the genus level. For example, age increased the relative abundance of Parabacteriodes, Oscillospira and Mucispirillum, which are associated with intestinal disorders and infections. CR reduced the abundance of these bacteria potentially exerting health benefits. The genus Parabacteriodes, which is significantly increased in old AL fed mice, are capable of producing toxic endotoxins, and some species of Parabacteriodes are strongly associated with increased inflammation and infections (Awadel-Kariem et al., 2010; Agudelo-Ochoa et al., 2020). CR significantly reduced the abundance of Parabacteriodes in old mice potentially reducing inflammation and susceptibility to infections. Similarly, age increased the abundance of the Oscillospira genus and Mucispirillum schaedleri both of which are associated with bowel diseases (Chen et al., 2020; Krych et al., 2015; Toit, 2019), and CR significantly reduced their abundance. On the other hand, age decreased the overall abundance of the Bacteriodes genus and Bacteriodes acidifaciens, and CR increased these in both the cecum and colon of C57BL/6JN mice. Microbes in the Bacteriodes genus are known to efficiently ferment carbohydrates producing fatty acids, which are a good fuel source for the intestine (Wexler et al., 2007). Bacteriodes acidifaciens has been shown to prevent obesity and improve insulin sensitivity (Yang et al., 2016). The improved insulin sensitivity observed in CR mice potentially could arise from the abundance of these health promoting bacteria. As discussed above, the Bacteriodes and Parabacteriodes genus also display a dual nature with some reports showing anti-inflammatory effects and some reports pro-inflammatory effects (Hiippala et al., 2020; Wexler et al., 2007). Therefore, the interpretation of the effect of changes in microbiome on the health of an organism needs to be done cautiously and more thorough functional analysis are warranted.

We are the first group to simultaneously compare the effect of life-long CR on the microbiome of two strains of mice (C57BL/6JN and B6D2F1) to determine if there were major strain differences with respect to the effect of CR on the microbiome. Different strains of mice have been reported to show differences in the microbiome (Laukens et al., 2015). However, Turturro et al (1999) showed that CR increased the lifespan of these two strains of mice to a similar extent (e.g., ∼25-35%). CR significantly altered the microbiome of old B6D2F1 mice and increased the diversity of the microbiome and the Bacteriodetes/ Firmicutes ratio in the old B6D2F1 mice as it did in the old C57BL/6JN mice. Similarly, old AL B6D2F1 mice showed an increase in the abundance in the Lachnospiraceae and Ruminocococcaeae families and the genus Parabacteriodes just as that observed in the C57Bl/6JN mice. CR increased the Rikenellaceae and S24-7 families, the genus Bacteriodes, and Bacteriodes acidifaciens in the B6D2F1 mice just as it did in the C57BL/6JN mice. The most notable difference between the C57BL/6JN and B6D2F1 microbial profile was in the abundance of Akkermansia muciniphila and the Lactobacillus genus. Akkermansia muciniphila was not identified in C57BL/6JN but was found to be significantly increased in the old B6D2F1 mice fed AL diet and this increase was reversed by CR. Akkermansia muciniphila is a mucin degrading bacteria under the phylum Verrucomicrobia and falls under the dual nature phenomenon of good versus bad microbes. Akkermansia muciniphila is generally considered to be a beneficial bacterium capable of conferring protection against obesity and inflammation (Xu et al., 2020); however, Akkermansia muciniphila has been reported to be abundant in the gut of patients with neurodegenerative diseases (Cirstea et al., 2018). The other major bacteria that was found to be different between C57BL/6JN and B6D2F1 mice is the Lactobacillus genus. Old AL B6D2F1 mice had a reduced abundance of Lactobacillus, and CR increased its abundance significantly. However, we did not observe a significant change in this genus in C57BL/6JN mice. The Lactobacillus genus is the most common bacteria used in probiotics to protect against pathogen invasion and inflammatory bowel disorders (Martin et al., 2013). Our Lactobacillus data in B6D2F1 are similar to that previously reported for C57BL/6 mice by Zhang et al. (2013) and Kok et al. (2018) showing that life-long CR resulted in an increase in the Lactobacillus genus. The difference in microbial types observed between studies again emphasizes the influence of environment and strain have on the gut microbiome (Ericcson et al., 2018; Laukens et al., 2015).

To gain a better understanding of the mechanism of how aging and CR affect the gut microbiome, we compared the effect of age and CR on the relative levels of microbes in both the cecum and colon. Most of the previous reports on aging studied only the colon microbiome, and this is the first study to compare the effects of age and CR on these two intestinal segments. We wanted to know if the age-related changes in the microbiome were limited to the colon or if changes were also seen in the cecum. Our data clearly show that significant changes in the microbiome occured in the cecum, e.g., ∼80% of the microbes in the colon of C57BL/6JN mice that changed significantly with age and reversed by CR were also significantly changed in the cecum. However, there were microbes in the cecum that CR altered and that were not altered in the colon and vice versa. In addition, the relative changes in the levels of microbes induced by CR also varied in the cecum and colon. For example, the magnitude of the age-related changes in Parabacteriodes and Ruminoccus were much greater in the cecum than colon. Parabacteriodes increased ∼7-fold in cecum and ∼2-fold in colon, and Ruminoccus was increased 100-fold in the cecum and ∼30-fold in the colon. Thus, our data show that changes in the microbiome with age and CR are initiated in the cecum, and these changes are further modified in the colon.

One of the major observations from this study was that 5 months of CR in adult, 9-month-old mice had only a small effect on the gut microbiome compared to life-long CR in old mice. While CR significantly altered the relative abundance of some microbes in the adult mice, the overall effect on the microbiome was minor compared to life-long CR in old mice, e.g., the PCA analysis (Figure 1) and the pattern of microbes (Figure 3) in adult mice fed AL or CR were similar for the cecum and colon. In addition, short-term CR had no significant effect on the diversity or the Bacteriodetes/ Firmicutes ratio in the adult mice while both were increased by CR in old mice (Figure 2). Fabbiano et al (2018) reported that short-term CR (40% restriction for 3 and 6 weeks) changed the microbiome in C57BL/6J mice; however, only a few weeks of CR in young mice was evaluated in this study. There was no comparison of these early changes to the effect of longer CR or long-term CR in old mice, which was used in our study.

Our data on the metabolome of fecal material paralleled the changes we observed with the microbiome. The effects of age and CR on the metabolome are broadly anti-correlated as one would predict from the microbiome data. For example, we observed that the age-related changes in the metabolome were attenuated by CR in C57BL/6JN mice and short-term CR had minimal effect on the fecal metabolome. Interestingly, the fecal metabolome was quite different in the old C57BL/6JN and B6D2F1 fed AL even though the microbiomes were similar.

One of the major questions that arises from studies on the effect of aging and CR on the microbiome is whether the changes in the microbiome play a role in the anti-aging actions of CR or if the physiological changes that arise from CR, e.g., the gastrointestinal (GI) system, are responsible for attenuating the age-related changes in the microbiome. To begin to answer this question, we studied the transcriptome of intestinal mucosal as a surrogate measure of intestinal status. Life-long CR in old mice had a dramatic effect on the transcriptome of intestinal mucosa, e.g., the transcripts of 250 genes were changed significantly (>=2.0-fold) by CR, and CR reversed the age-related changes in over 75% (189 genes) of the transcripts that changed significantly with age. Based on the large literature showing that life-long CR has a major effect of the transcriptome in various tissues (Barger et al., 2017), we were not surprised to observe a major effect of long-term CR on the transcriptome of the GI system, even though this is the first study to our knowledge to evaluate the effect of life-long CR on intestinal mucosa. However, we were surprised to find that short-term CR in adult mice also had a major effect on the transcriptome, e.g., PCA analysis of the transcripts showed that CR in both adult and old mice resulted in a significant change in the transcriptome. This observation was in contrast to what we observed for the microbiome and the fecal metabolome, where we observed only minor effects of 5 months of CR in adult mice. Thus, our transcriptome data show that CR has an impact on the colon before any major changes in the microbiome occur and agree with previous studies reporting that short-term CR has a significant impact on the gastrointestinal system. For example, early studies from Holt’s laboratory showed that 10 to 16 weeks of CR reduced epithelial cell proliferation in the rat colon (Steinbach et al., 1993) and rectal cell proliferation in humans (Steinbach et al., 1994). Albanes et al (1990) showed that 3 weeks of CR reduced DNA synthesis in the colon of young rats, which resulted in a decrease in the number of crypts and the total number of colonic mucosal cells dividing at any given time. In addition, 6-12 weeks of CR has been shown to have a significant impact on intestinal stem cell function in mice (Yilmaz et al., 2012; Yousefi et al., 2018), and we found that one-month of CR altered the expression of over 5000 genes (fold change >1.25) in intestinal mucosa from mice (Unnikrishnan et al., 2017). Thus, our data and previous data on the impact of CR on the GI-system indicate that the initial impact of CR occurs on the physiological status of the GI-system, suggesting that the subsequent changes in the microbiome occur because of these physiological changes.

In summary, our data demonstrate that life-long CR has a dramatic effect on the microbiome in old mice, which was similar in two strains of mice. The changes in the microbiome that occur with age and CR are initiated in the cecum and modified as the fecal material progresses to the colon. Importantly, our study gives us the first insight into the role the microbiome and the physiological status of the gastrointestinal system play in aging and the anti-aging actions of CR. Our data lead us to propose that the primary impact of CR is on the physiological status of the host’s gastrointestinal system, maintaining it in a more youthful state, which in turn maintains a more diverse and ‘youthful’ microbiome.

## Materials and methods

### Animals

Male C57BL/6JN and B6D2F1 mice fed either AL or CR were obtained from the aging colony maintained by NIA. After receiving the mice from NIA, animals were housed at the animal facility and maintained under SPF conditions and individually housed in a HEPA barrier environment at the University of Oklahoma Health Sciences Center for at 4-8 weeks before being used in the following experiments. Mice fed *ad libitum* were fed irradiated NIH-31 mouse/rat diet (Teklad, Envigo), and the CR mice were fed the same diet fortified for micronutrients. CR was initiated by NIA at 14 weeks of age, at a level of 10% restriction, increased to 25% restriction at 15 weeks, and to 40% restriction at 16 weeks of age. Adult (9 months of age) fed AL or CR and old mice (24 months of age) fed AL or CR were fasted overnight, and samples were collected during sacrifice (n=8-10/group). Mice were euthanized by decapitation and fecal material from cecum and colon and colon mucosa were harvested, snap frozen, and stored at −80°C until analyzed. Microbiome, metabolome and transcriptome analysis were done in the same animals. All animal experiments were performed according to protocols approved by the Institutional Animal Care and Use Committee.

### Microbiome Analysis

#### DNA extraction

Fecal DNA was extracted from luminal fecal samples of cecum and colon using PowerFecal kits (Qiagen) according to the manufacturer’s instructions, with the exception that, rather than performing the initial homogenization of samples using the vortex adapter described in the protocol, samples are homogenized in the provided bead tubes using a TissueLyser II (Qiagen, Venlo, Netherlands) for three minutes at 30/sec, before proceeding according to the protocol and eluting in 100 µL of elution buffer (Qiagen). DNA yields are quantified via fluorometry (Qubit 2.0, Invitrogen, Carlsbad, CA) using quant-iT BR dsDNA reagent kits (Invitrogen).

#### 16S rRNA library preparation and sequencing

Extracted fecal DNA was processed at the University of Missouri DNA Core Facility. Bacterial 16S rDNA amplicons are constructed via amplification of the V4 region of the 16S rRNA gene with universal primers (U515F/806R) previously developed against the V4 region, flanked by Illumina standard adapter sequences. Oligonucleotide sequences are available at proBase (Greuter et al., 2016). Dual-indexed forward and reverse primers were used in all reactions. PCR was performed in 50 µL reactions containing 100 ng metagenomic DNA, primers (0.2 µM each), dNTPs (200 µM each), and Phusion high-fidelity DNA polymerase (1U). Amplification parameters were 98°C^(3:00)^ + [98°C^(0:15)^ + 50°C^(0:30)^ + 72°C^(0:30)^] × 25 cycles +72°C^(7:00)^. Amplicon pools (5 µL/reaction) were combined, thoroughly mixed, and then purified by addition of Axygen Axyprep MagPCR clean-up beads to an equal volume of 50 µL of amplicons and incubated for 15 minutes at room temperature. Products were then washed multiple times with 80% ethanol, and the dried pellet was resuspended in 32.5 µL EB buffer, incubated for two minutes at room temperature, and then placed on the magnetic stand for five minutes. The final amplicon pool was evaluated using the Advanced Analytical Fragment Analyzer automated electrophoresis system, quantified using quant-iT HS dsDNA reagent kits, and diluted according to Illumina’s standard protocol for sequencing on the MiSeq instrument.

#### Informatics analysis

Read merging, clustering, and annotation of DNA sequences was performed at the University of Missouri Informatics Research Core Facility. Paired DNA sequences were merged using FLASH software (Magoč et al., 2011), and removed if found to be far from the expected length of 292 bases after trimming for base quality of 31. Cutadapt (https://github.com/marcelm/cutadapt) was used to remove the primers at both ends of the contig and cull contigs that did not contain both primers. The usearch (Edgar, 2010) fastq_filter command (http://drive5.com/usearch/manual/cmd_fastq_filter.html) was used for quality trimming of contigs and rejecting those for which the expected number of errors is greater than 0.5. All contigs were trimmed to 248 bases and shorter contigs are removed. The Qiime 1.9 (Caporaso et al., 2010) command split_libraries_fastq.py was used to demultiplex the samples. The outputs for all samples were combined into a single file for clustering. The uparse method (http://www.drive5.com/uparse/) was used to both cluster contigs with 97% identity and remove chimeras. Taxonomy was assigned to selected OTUs using BLAST (Altschul et al., 1990) against the SILVA database v132 (Quast et al., 2012) of 16S rRNA sequences and taxonomy.

#### Statistical Analysis

Relative abundances were performed through Qiime 1.9, and all the bar graphs were generated using Microsoft Excel software (Microsoft, Seattle, WA). Principal components analysis was performed using the scikit-learn Python package (Pedregosa et al., 2011), and OLS regression was performed using the statsmodels package (Seabold et al., 2010).

### Metabolome Analysis

Metabolomics analysis of the fecal pellets from the colon was performed in collaboration with the Nathan Shock Center of Excellent in Basic Biology of Aging at the University of Washington. The targeted liquid chromatography mass spectrometry (LC-MS/MS) data were collected using a standard protocol developed by the Northwest Metabolomics Research Center (NW-MRC) that has been used in a number of studies (Zhu et al., 2014; Carroll et al., 2015; Sperber et al., 2015; Rabinowitz et al., 2017; Parent et al., 2017; Li et al., 2018; Shao et al., 2018). Briefly, the LC-MS/MS experiments were performed on a Shimadzu Nexera LC-20ADXR (Shimadzu, Kyoto, Japan) AB Sciex Triple Quad 6500+ MS (AB Sciex, Toronto, Canada) system equipped with a PAL HTC-xt autosampler (CTC Analytics, Zwingen, Switzerland). Each sample was injected twice, 10 µL for analysis using negative ionization mode and 5 µL for analysis using positive ionization mode. Both chromatographic separations were performed using hydrophilic interaction chromatography (HILIC) on a Waters XBridge BEH Amide column (150 × mm, 2.5 µm particle size, Waters Corporation, Milford, MA). The flow rate was 0.3 mL/min. The mobile phase was composed of Solvents A (10 mM ammonium acetate in 95% H_2_O/ 3% acetonitrile/ 2% methanol + 0.2% acetic acid) and B (10 mM ammonium acetate in 93% acetonitrile/ 5% H_2_O / 2% methanol+ 0.2% acetic acid). After the initial 3 min isocratic elution of 95% B, the percentage of Solvent B was decreased linearly to 50% at t=8 min. The composition of Solvent B was maintained at 50% for 4 min (t=12 min), and then the percentage of B was gradually increased to 95%, to prepare for the next injection. The metabolite identities were confirmed by spiking the pooled serum sample used for method development with mixtures of standard compounds. Isotope labeled metabolite standards were spiked into the samples at different times during the sample preparation to monitor sample processing and sample injections. A laboratory quality control sample and pooled sample QC sample were measured once for every 10 biological samples to monitor data quality and provide consistent metabolite signals to normalize for any instrument drift. The extracted MRM peaks were integrated using MultiQuant 3.0.2 software (AB Sciex, Toronto, Canada).

#### Statistical Analysis

Analysis of the targeted metabolomics data with 36 samples and 216 metabolites was performed using R (version 3.4.2). Metabolite abundance was log_2_-transformed and median normalized prior to imputation. All the metabolites with ≥ 5% missingness were excluded, and a total of 98 metabolites were included in the imputation step. We imputed the remaining missing values using the K-nearest neighbors imputation method implemented in the R impute package (Troyanskaya et al., 2001). We fit a weighted linear model to the normalized and imputed metabolomic data using the Bioconductor limma package to test the group differences (Ritchie et al. 2015). The limma package uses empirical Bayes moderated statistics, which improves power by ‘borrowing strength’ between metabolites to moderate the residual variance (G. Smyth 2004). By using a weighted linear regression, we can smoothly up or down-weight individual samples, based on similarity to other similar sample types (Ritchie et al., 2006). This allows us to keep all samples in the analysis, and it also minimizes the need to make decisions about removing possible outlier samples from consideration. We selected metabolites that have a significant difference with a 5% false discovery rate (FDR) using the Benjamini-Hochberg method (Benjamini and Hochberg 1995).

### Transcriptome Analysis

Transcriptome analysis was done using strand-specific RNA-Seq technology for RNA isolated from the mucosa from the colon of adult and old mice fed AL or CR using the genomic sequencing facility at OMRF. RNA was isolated from the colon mucosa using the RNeasy kit from Qiagen (Germantown MD, USA). RNA integrity was checked using the Bioanalyzer (Agilent) and only samples with RNA integrity numbers >8 were used in the RNA-Seq. RNA was depleted of ribosomal RNA (Ribozero, Illumina) and used for generation of stranded RNA sequencing libraries, whereby the orientation and originating DNA strand of the RNA are maintained (Illumina Stranded RNA-Seq). Each library was uniquely indexed and then sized and quantified by capillary electrophoresis (TapeStation, Agilent). Libraries were sequenced in a paired-end 75 fashion on an Illumina HiSeq 2000 in rapid run mode. An average of 81.1±9.5 million reads were generated for each library.

Raw FASTQ files were imported into Strand NGS software (Strand Life Sciences, Version 3.4, Build 239479) for trimming and alignment. All the statistical analysis and figure generation on transcriptome data were performed using Strand NGS software. Reads with average base Q score of <30 were discarded, and 1 base from 3’ and 5’, and adaptor sequences were eliminated. Reads were then aligned to mouse, build mm10 (UCSC) in an orientation-specific fashion. The transcript annotation used was UCSC transcripts (2018.02.25). Reads that aligned normally (in the appropriate direction) were retained. Reads were further filtered on read quality metrics using the following cut-offs: N’s allowed in read <=0, quality threshold >= 30, number of multiple matches allowed <= 1. Duplicate reads were also excluded. Data were then normalized to the AL group and quantification was performed with the DESeq algorithm (Anders and Huber, 2010). Transcripts that had a raw read count value >20 in 100% of the samples and in at least one of the conditions were considered expressed at a level sufficient for quantitation and retained. Statistically significant differentially expressed genes with P (Corr) cut off < 0.05, FC cut off 2.0 were determined using One-way ANOVA (Multiple testing correction: Benjamini-Hochberg, Post hoc test: Student Newman Keuls).

## Acknowledgements

The research was supported by the following NIH grants: R01AG045693 (AU and AR), R01AG049494 (DP), R21AG062894 (AR), KO1AG 056655-01A1 (AU), S10 OD021562-01 (DR) and the Nathan Shock Centers of Oklahoma, P30 AG050911 (AR and JW) and University of Washington, P30 AG013280 (DP). DP was also supported in part by R01AG049494. Additional support came from the Oklahoma Center for Advancement of Science and Technology HR17-098 (AU), The Oklahoma Center for Adult Stem Cell Research (AR), the American Federation of Aging Research 17132 (AU), and a Senior Career Research Award (AR) from the Department of Veterans Affairs.

