## Supporting File for "Calorie Restriction Prevents Age-Related Changes in the Intestinal Microbiota"

**Figure 1S: Relative abundance of all the microbes in the cecum and colon of C57BL/6J mice**

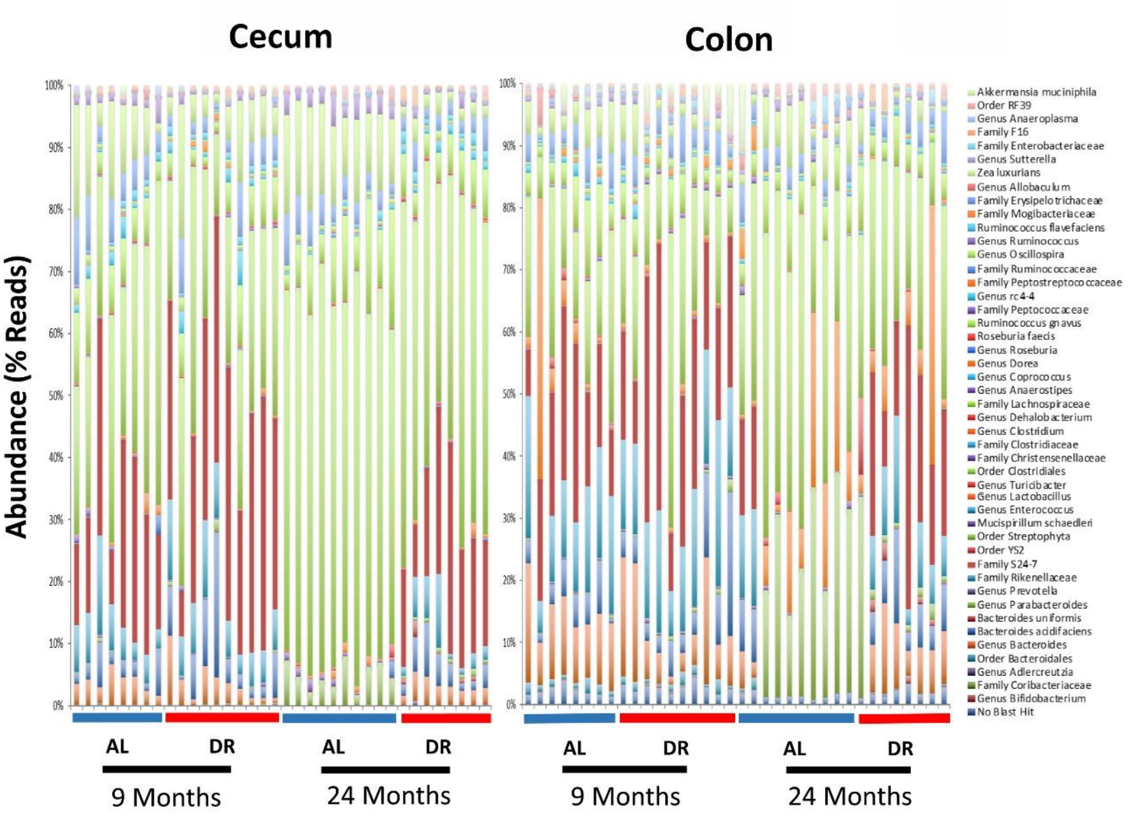

**Figure 2S: Relative abundance of all the microbes in the cecum and colon of the B6D2F1 mice**

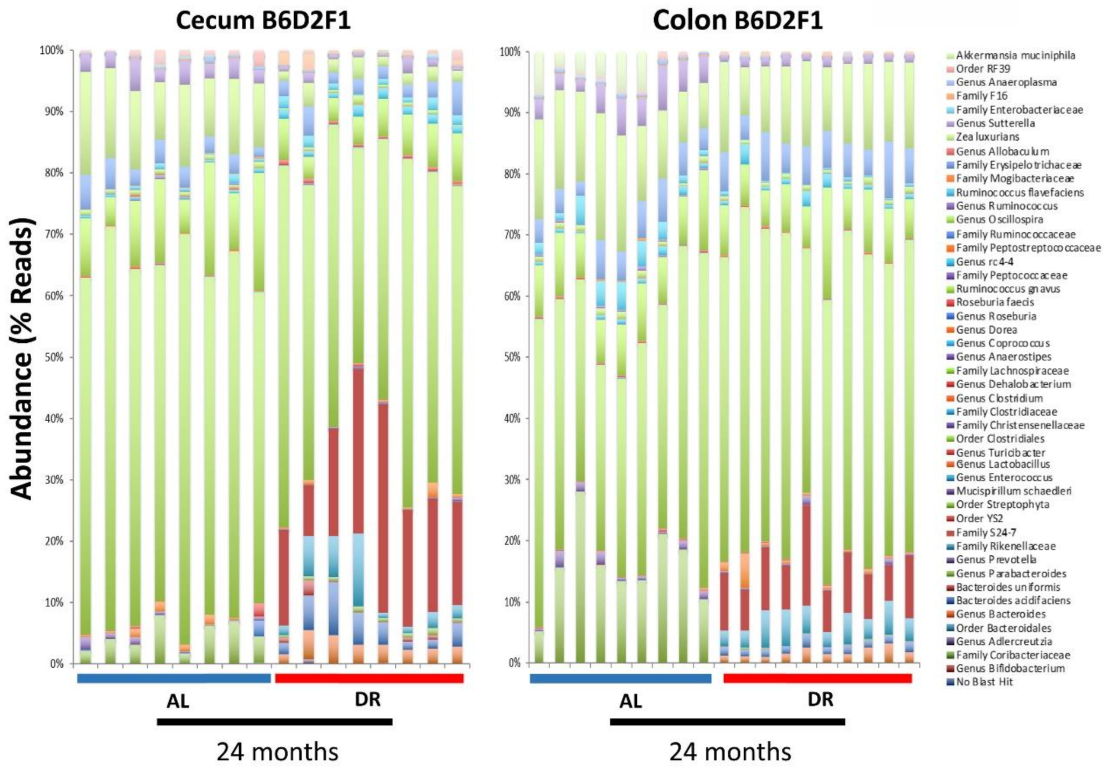

**Table 1S: Microbiota Composition of the Cecum from Adult and Old C57BL/6 Mice Fed *Ad libitum* (AL) or Caloric Restriction (CR)**

| | Adult AL | Adult CR | Old AL | Old CR |
|---|---|---|---|---|
| **Species** | | | | |
| Akkermansia muciniphila | 0.43 ± 0.36 | 3.25 ± 1.16$^{a}$ | 0.0 ± 0.0 | 0.01 ± 0.01$^{d}$ |
| Bacteroides acidifaciens | 4.33 ± 0.72 | 8.1 ± 2.08 | 0.0 ± 0.0$^{b}$ | 7.15 ± 1.56$^{c}$ |
| Bacteroides uniformis | 0.01 ± 0.0 | 0.0 ± 0.0 | 0.0 ± 0.0$^{b}$ | 1.12 ± 0.39$^{cd}$ |
| Eubacterium cylindroides | 0.01 ± 0.01 | 0.55 ± 0.16$^{a}$ | 0.0 ± 0.0 | 0.0 ± 0.0$^{d}$ |
| Mucispirillum schaedleri | 0.16 ± 0.02 | 0.11 ± 0.03 | 0.45 ± 0.13$^{b}$ | 0.04 ± 0.01$^{c}$ |
| Ruminococcus gnavus | 0.7 ± 0.12 | 0.49 ± 0.11 | 0.45 ± 0.04 | 0.26 ± 0.05$^{c}$ |
| **Genus** | | | | |
| Genus Allobaculum | 0.88 ± 0.61 | 0.55 ± 0.25 | 0.0 ± 0.0 | 0.0 ± 0.0 |
| Genus Anaeroplasma | 0.52 ± 0.16 | 0.0 ± 0.0$^{a}$ | 0.04 ± 0.02$^{b}$ | 0.0 ± 0.0 |
| Genus Bacteroides | 13.23 ± 1.51 | 6.47 ± 1.74$^{a}$ | 0.0 ± 0.0$^{b}$ | 8.75 ± 1.26$^{c}$ |
| Genus Coprococcus | 1.13 ± 0.24 | 0.41 ± 0.05$^{a}$ | 0.37 ± 0.06$^{b}$ | 0.55 ± 0.12 |
| Genus Dehalobacterium | 0.19 ± 0.03 | 0.17 ± 0.03 | 0.11 ± 0.02$^{b}$ | 0.13 ± 0.03 |
| Genus Dorea | 0.38 ± 0.16 | 1.66 ± 0.52$^{a}$ | 0.05 ± 0.01 | 0.14 ± 0.04$^{cd}$ |
| Genus Lactobacillus | 6.26 ± 4.47 | 0.9 ± 0.1 | 12.46 ± 3.24 | 9.27 ± 5.64 |
| Genus Oscillospira | 5.14 ± 0.75 | 1.61 ± 0.22$^{a}$ | 6.36 ± 0.95 | 1.08 ± 0.41$^{c}$ |
| Genus Parabacteroides | 0.2 ± 0.05 | 0.56 ± 0.19 | 25.99 ± 2.89$^{b}$ | 1.45 ± 0.22$^{cd}$ |
| Genus Prevotella | 0.26 ± 0.05 | 0.55 ± 0.09$^{a}$ | 0.0 ± 0.0$^{b}$ | 0.0 ± 0.0$^{d}$ |
| Genus Ruminococcus | 0.71 ± 0.13 | 0.37 ± 0.1$^{a}$ | 1.07 ± 0.13 | 0.36 ± 0.15$^{c}$ |
| Genus Turicibacter | 0.52 ± 0.25 | 0.0 ± 0.0 | 1.85 ± 1.36 | 0.17 ± 0.1 |
| Genus rc4-4 | 0.47 ± 0.07 | 0.45 ± 0.05 | 0.29 ± 0.11 | 0.43 ± 0.08 |
| **Family** | | | | |
| Family Christensenellaceae | 0.2 ± 0.04 | 0.79 ± 0.09$^{a}$ | 0.04 ± 0.01$^{b}$ | 0.08 ± 0.02$^{d}$ |
| Family Clostridiaceae | 0.13 ± 0.03 | 0.09 ± 0.01 | 0.13 ± 0.06 | 0.1 ± 0.03 |
| Family Enterobacteriaceae | 0.04 ± 0.02 | 0.04 ± 0.02 | 0.91 ± 0.48 | 0.43 ± 0.3 |
| Family Erysipelotrichaceae | 0.47 ± 0.11 | 0.33 ± 0.12 | 0.98 ± 0.28 | 0.43 ± 0.08 |
| Family F16 | 0.26 ± 0.06 | 0.61 ± 0.1$^{a}$ | 0.0 ± 0.0$^{b}$ | 1.17 ± 0.55$^{c}$ |
| Family Lachnospiraceae | 6.05 ± 1.18 | 3.78 ± 0.63 | 6.72 ± 0.99 | 4.0 ± 0.78 |
| Family Mogibacteriaceae | 0.1 ± 0.01 | 0.09 ± 0.01 | 0.35 ± 0.08$^{b}$ | 0.09 ± 0.02$^{c}$ |
| Family Rikenellaceae | 14.93 ± 1.61 | 16.81 ± 1.66 | 0.0 ± 0.0$^{b}$ | 9.2 ± 1.79$^{cd}$ |
| Family Ruminococcaceae | 4.01 ± 0.51 | 3.9 ± 0.66 | 2.62 ± 0.29$^{b}$ | 3.91 ± 0.92 |
| Family S24-7 | 17.83 ± 2.37 | 24.17 ± 3.53 | 0.0 ± 0.0$^{b}$ | 23.1 ± 4.78$^{c}$ |
| **Order** | | | | |
| Order Bacteroidales | 1.42 ± 0.2 | 1.21 ± 0.27 | 0.0 ± 0.0$^{b}$ | 0.12 ± 0.03$^{cd}$ |
| Order Clostridiales | 18.41 ± 2.37 | 20.93 ± 4.17 | 37.09 ± 4.01$^{b}$ | 25.8 ± 3.16 |
| Order RF32 | 0.16 ± 0.05 | 0.28 ± 0.12 | 0.0 ± 0.0$^{b}$ | 0.0 ± 0.0 |
| Order RF39 | 0.24 ± 0.06 | 0.55 ± 0.11$^{a}$ | 1.67 ± 0.21$^{b}$ | 0.67 ± 0.22$^{c}$ |
| Order YS2 | 0.22 ± 0.1 | 0.23 ± 0.07 | 0.0 ± 0.0 | 0.0 ± 0.0$^{d}$ |

Each value represents the mean ± SEM of data generated from 8 to 10 mice per group. Significant differences between groups are shown for an FDR <0.05: a = Significant difference between adult AL and young CR; b = Significant difference between adult AL and old AL; c = Significant difference between old AL and old CR; d = Significant difference between adult CR and old CR.

**Table 2S: Microbiota Composition of Colon from Adult and Old C57BL/6 Mice Fed *Ad Libitum* (AL) or Caloric Restriction (CR)**

| | Adult AL | Adult CR | Old AL | Old CR |
|---|---|---|---|---|
| **Species** | | | | |
| Bacteroides acidifaciens | 2.75 ± 0.84 | 6.71 ± 2.15 | 0.0 ± 0.0[b] | 3.97 ± 0.89[c] |
| Bacteroides uniformis | 0.0 ± 0.0 | 0.0 ± 0.0 | 0.0 ± 0.0 | 0.67 ± 0.29[cd] |
| Mucispirillum schaedleri | 0.47 ± 0.18 | 0.19 ± 0.09 | 0.87 ± 0.2 | 0.24 ± 0.07[c] |
| Ruminococcus gnavus | 0.79 ± 0.27 | 0.65 ± 0.17 | 0.78 ± 0.06 | 0.51 ± 0.13 |
| **Genus** | | | | |
| Genus Anaeroplasma | 0.37 ± 0.18 | 0.1 ± 0.08 | 0.01 ± 0.0 | 0.0 ± 0.0 |
| Genus Bacteroides | 2.95 ± 0.67 | 3.54 ± 1.03 | 0.0 ± 0.0[b] | 3.02 ± 0.41[c] |
| Genus Coprococcus | 1.28 ± 0.37 | 1.72 ± 0.54 | 0.44 ± 0.04[b] | 1.34 ± 0.14[c] |
| Genus Dehalobacterium | 0.22 ± 0.05 | 0.39 ± 0.05[a] | 0.24 ± 0.03 | 0.54 ± 0.06[c] |
| Genus Dorea | 0.08 ± 0.04 | 0.56 ± 0.17[a] | 0.02 ± 0.01 | 0.07 ± 0.02[cd] |
| Genus Lactobacillus | 0.65 ± 0.34 | 0.26 ± 0.1 | 0.7 ± 0.2 | 0.57 ± 0.27 |
| Genus Oscillospira | 8.18 ± 2.13 | 7.14 ± 1.91 | 13.13 ± 0.83[b] | 2.36 ± 0.31[cd] |
| Genus Parabacteroides | 0.13 ± 0.08 | 0.41 ± 0.21 | 4.79 ± 0.7[b] | 0.73 ± 0.08[c] |
| Genus Ruminococcus | 1.49 ± 0.41 | 0.79 ± 0.17 | 3.02 ± 0.34[b] | 1.06 ± 0.29[c] |
| Genus Turicibacter | 0.04 ± 0.03 | 0.01 ± 0.0 | 0.38 ± 0.22 | 0.09 ± 0.06 |
| Genus rc4-4 | 0.52 ± 0.19 | 0.97 ± 0.12 | 0.44 ± 0.16 | 1.3 ± 0.24[c] |
| **Family** | | | | |
| Family Christensenellaceae | 0.11 ± 0.03 | 0.25 ± 0.05[a] | 0.01 ± 0.0[b] | 0.09 ± 0.02[cd] |
| Family Erysipelotrichaceae | 0.08 ± 0.04 | 0.21 ± 0.09 | 0.3 ± 0.1 | 0.21 ± 0.05 |
| Family F16 | 0.25 ± 0.08 | 0.48 ± 0.08 | 0.0 ± 0.0[b] | 0.85 ± 0.34[c] |
| Family Lachnospiraceae | 5.8 ± 1.35 | 5.15 ± 0.9 | 10.04 ± 1.84 | 5.82 ± 0.58 |
| Family Rikenellaceae | 5.52 ± 1.51 | 7.57 ± 1.02 | 0.01 ± 0.0[b] | 4.17 ± 1.37[c] |
| Family Ruminococcaceae | 5.74 ± 1.16 | 3.96 ± 0.9 | 4.28 ± 0.64 | 3.0 ± 0.5 |
| Family S24-7 | 26.57 ± 4.76 | 31.62 ± 3.34 | 0.02 ± 0.01[b] | 19.72 ± 2.75[cd] |
| Family Sphingobacteriaceae | 0.32 ± 0.31 | 0.02 ± 0.0 | 0.29 ± 0.27 | 0.02 ± 0.0 |
| **Order** | | | | |
| Order Clostridiales | 35.52 ± 3.77 | 27.0 ± 2.59 | 59.33 ± 1.49[b] | 49.17 ± 2.67[cd] |
| Order RF39 | 0.16 ± 0.05 | 0.31 ± 0.07 | 0.89 ± 0.18[b] | 0.48 ± 0.18 |

Each value represents the mean ± SEM of data generated from 8 to 10 mice per group. Significant differences between groups are shown for an FDR <0.05: a = significant difference between adult AL and adult CR; b = significant difference between adult AL and old AL; c = Significant difference between old AL and old CR; d = Significant difference between adult CR and old CR.

**Table 3S: Microbiota Composition of Cecum and Colon from Old B6D2F1 Mice Fed *Ad Libitum* (AL) or Caloric Restriction (CR)**

| | Cecum | | Colon | |
|---|---|---|---|---|
| | **Old AL** | **Old CR** | **Old AL** | **Old CR** |
| **Species** | | | | |
| Akkermansia muciniphila | 7.11 ± 2.01 | 0.0 ± 0.0a | 3.06 ± 0.94 | 0.0 ± 0.0[a] |
| Bacteroides acidifaciens | 0.0 ± 0.0 | 5.78 ± 0.43[a] | 0.67 ± 0.67 | 1.4 ± 0.1 |
| Mucispirillum schaedleri | 0.2 ± 0.13 | 0.03 ± 0.01 | 1.1 ± 0.28 | 0.47 ± 0.15 |
| Ruminococcus gnavus | 0.52 ± 0.1 | 0.38 ± 0.06 | 0.79 ± 0.11 | 0.75 ± 0.06 |
| **Genus** | | | | |
| Genus Adlercreutzia | 0.09 ± 0.02 | 0.17 ± 0.02[a] | ND | ND |
| Genus Anaeroplasma | 0.17 ± 0.08 | 0.09 ± 0.09 | ND | ND |
| Genus Anaerostipes | 0.26 ± 0.03 | 0.0 ± 0.0[a] | ND | ND |
| Genus Bacteroides | 0.0 ± 0.0 | 4.67 ± 0.63[a] | 0.0 ± 0.0 | 1.67 ± 0.24[a] |
| Genus Coprococcus | 0.45 ± 0.07 | 0.75 ± 0.14 | 0.74 ± 0.14 | 1.29 ± 0.32 |
| Genus Dehalobacterium | 0.26 ± 0.04 | 0.17 ± 0.01[a] | 0.28 ± 0.04 | 0.27 ± 0.02 |
| Genus Lactobacillus | 0.1 ± 0.04 | 5.09 ± 0.67[a] | 0.06 ± 0.04 | 1.05 ± 0.52 |
| Genus Oscillospira | 6.78 ± 0.36 | 5.16 ± 0.61[a] | 12.68 ± 1.97 | 12.47 ± 0.68 |
| Genus Parabacteroides | 30.72 ± 2.64 | 0.21 ± 0.05[a] | 14.18 ± 2.51 | 0.11 ± 0.03[a] |
| Genus Prevotella | 0.0 ± 0.0 | 0.51 ± 0.14[a] | ND | ND |
| Genus Ruminococcus | 2.33 ± 0.38 | 0.56 ± 0.07[a] | 3.97 ± 0.66 | 1.46 ± 0.1[a] |
| Genus Turicibacter | 0.6 ± 0.21 | 1.17 ± 0.18 | 0.18 ± 0.04 | 0.14 ± 0.02 |
| Genus rc4-4 | 5.01 ± 1.09 | 0.8 ± 0.15[a] | 2.54 ± 0.55 | 0.32 ± 0.06[a] |
| **Family** | | | | |
| Family Erysipelotrichaceae | 0.92 ± 0.18 | 0.06 ± 0.02[a] | 0.45 ± 0.09 | 0.04 ± 0.01[a] |
| Family F16 | 0.0 ± 0.0 | 1.05 ± 0.22[a] | 0.0 ± 0.0 | 0.27 ± 0.03[a] |
| Family Lachnospiraceae | 6.92 ± 0.67 | 3.83 ± 0.49[a] | 8.64 ± 0.62 | 8.55 ± 1.15 |
| Family Mogibacteriaceae | 0.18 ± 0.02 | 0.09 ± 0.01[a] | 0.16 ± 0.03 | 0.06 ± 0.01[a] |
| Family Peptostreptococcaceae | 0.29 ± 0.15 | 0.01 ± 0.01 | 0.29 ± 0.07 | 0.07 ± 0.01[a] |
| Family Rikenellaceae | 0.0 ± 0.0 | 7.18 ± 0.72[a] | 0.67 ± 0.67 | 4.19 ± 0.46[a] |
| Family Ruminococcaceae | 4.03 ± 0.57 | 2.89 ± 0.39 | 5.76 ± 0.96 | 6.04 ± 0.51 |
| Family S24-7 | 0.02 ± 0.0 | 33.76 ± 1.99[a] | 2.68 ± 2.67 | 9.03 ± 0.97[a] |
| **Order** | | | | |
| Order Clostridiales | 32.29 ± 3.22 | 24.95 ± 1.58 | 40.66 ± 2.57 | 50.13 ± 1.43[a] |
| Order RF39 | 0.71 ± 0.13 | 0.63 ± 0.11 | 0.43 ± 0.13 | 0.24 ± 0.03 |

Each value represents the mean ± SEM of data generated from 8 to 10 mice per group. Significant differences between groups are shown for an FDR <0.05: a = significant difference between AL and CR mice. ND = Not detected.

**Table 4S: Levels of Fecal Metabolites from C57BL/6 and B6D2F1 Mice Fed *Ad Libitum* (AL) or Caloric Restriction (CR)**

| | C57BL/6 | | | | B6D2F1 | |
|---|---|---|---|---|---|---|
| **Metabolites** | **Adult AL** | **Adult CR** | **Old AL** | **Old CR** | **Old AL** | **Old CR** |
| 12-HETE (319.2 / 179.0) | 14.53 ± 0.19 | 15 ± 0.2 | 14.43 ± 0.21 | 14.9 ± 0.19 | 14.89 ± 0.25 | 14.91 ± 0.17 |
| 13-HODE (295.1 / 195.0) | 22.2 ± 0.35 | 22.3 ± 0.34 | 23.29 ± 0.29 | 23.1 ± 0.14 | 21.42 ± 0.27 | 21.23 ± 0.54 |
| 2-Hydroxyglutarate (147.0 / 129.0) | 20.88 ± 0.16 | 21.1 ± 0.28 | 21.13 ± 0.28 | 20.6 ± 0.08 | 21.84 ± 0.24 | 21.75 ± 0.46 |
| 2-Hydroxyisovaleric Acid (117.0 / 71.0) | 21.65 ± 0.28 | 22.1 ± 0.33 | 22.36 ± 0.73 | 22.5 ± 0.15 | 20.62 ± 0.37 | 21.36 ± 0.21 |
| 3HBA (103.0 / 59.0) | 19.32 ± 0.11 | 20.1 ± 0.2 | 18.98 ± 0.23a | 19.1 ± 0.26 | 19.55 ± 0.33 | 19.60 ± 0.41 |
| 3-Hydroxykynurenine (225.1 / 208.1) | 17.74 ± 0.29 | 18 ± 0.38 | 18.22 ± 0.13 | 18.7 ± 0.12 | 17.76 ± 0.16 | 18.38 ± 0.22 |
| 4-Hydroxybutyrate (105.0 / 77.0) | 21.56 ± 0.22 | 22.4 ± 0.15 | 22.15 ± 0.39 | 22.4 ± 0.29 | 22.34 ± 0.42 | 21.88 ± 0.38 |
| 4-Pyridoxic acid (182.1 / 138.0) | 22.55 ± 0.2 | 22.9 ± 0.21 | 22.44 ± 0.26 | 22.6 ± 0.15 | 22.31 ± 0.46 | 22.87 ± 0.22 |
| 5-Aminovaleric Acid (118.0 / 101.0) | 20.78 ± 0.5 | 20.5 ± 0.86 | 20.41 ± 0.64 | 19.2 ± 0.51 | 19.93 ± 0.79 | 19.06 ± 0.58 |
| Acetylcarnitine (204.1 / 85.0) | 17.88 ± 0.58 | 16.5 ± 0.69 | 17.37 ± 0.56 | 17.8 ± 0.29 | 17.13 ± 0.25 | 17.06 ± 0.88 |
| Adenine (134.0 / 107.0) | 24.36 ± 0.51 | 26.1 ± 0.35 | 21.68 ± 0.37a | 23.4 ± 0.61b | 22.17 ± 0.29 | 25.46 ± 0.25b |
| Adenosine (268.2 / 136.1) | 21.72 ± 0.18 | 21.5 ± 0.32 | 18.07 ± 0.31a | 20.2 ± 0.47b | 20.77 ± 0.19 | 20.21 ± 0.55 |
| Adipic Acid (144.9 / 83.0) | 16.32 ± 0.21 | 16.9 ± 0.33 | 16.34 ± 0.1 | 16.5 ± 0.15 | 16.17 ± 0.24 | 15.92 ± 0.26 |
| Alanine (90.0 / 44.0 (2)) | 20.29 ± 0.2 | 20.3 ± 0.23 | 21.01 ± 0.18a | 20.5 ± 0.22 | 21.85 ± 0.04 | 20.54 ± 0.11b |
| Allantoin (157.0 / 114.0) | 17.07 ± 0.26 | 16.2 ± 0.26 | 16.88 ± 0.26 | 16.5 ± 0.21 | 17.13 ± 0.19 | 16.40 ± 0.21b |

| | | | | | | |
|---|---|---|---|---|---|---|
| Alpha-Ketoglutaric Acid (145.0 / 101.0) | 21.01 ± 0.24 | 21.6 ± 0.23 | 21.17 ± 0.32 | 21 ± 0.13 | 22.92 ± 0.25 | 22.48 ± 0.68 |
| AMP (346.1 / 79.0) | 17.08 ± 0.53 | 17 ± 0.38 | 16.83 ± 0.27 | 15.9 ± 0.31 | 16.20 ± 0.52 | 17.34 ± 0.58b |
| Arachidonate (303.3 / 59.0) | 22.17 ± 0.38 | 23.9 ± 0.14 | 22.84 ± 0.22 | 23.4 ± 0.07 | 22.64 ± 0.19 | 22.63 ± 0.24 |
| Arginine (175.0 / 70.0) | 23.53 ± 0.19 | 21.7 ± 0.26 | 25.04 ± 0.14a | 23.3 ± 0.23b | 25.56 ± 0.15 | 22.93 ± 0.24b |
| Asparagine (133.0 / 74.0) | 17.58 ± 0.38 | 15.4 ± 0.35 | 19.25 ± 0.14a | 16.4 ± 0.24b | 20.34 ± 0.06 | 17.36 ± 0.34b |
| Aspartic Acid (134.0 / 74.0) | 20.02 ± 0.33 | 20.8 ± 0.28 | 20.84 ± 0.28a | 19.4 ± 0.24b | 21.91 ± 0.13 | 20.67 ± 0.14b |
| Azelaic Acid (187.0 / 125.0) | 24.35 ± 0.32 | 26.3 ± 0.33 | 24.93 ± 0.37 | 25.8 ± 0.27b | 23.71 ± 0.12 | 23.24 ± 0.37 |
| Betaine (118.0 / 58.0) | 22.24 ± 0.32 | 21.9 ± 0.19 | 22.21 ± 0.36 | 21.6 ± 0.13 | 21.52 ± 0.23 | 21.90 ± 0.18 |
| Cadaverine (103.0 / 86.0) | 15.86 ± 0.21 | 16.1 ± 0.3 | 15.73 ± 0.08 | 15.5 ± 0.18 | 15.96 ± 0.09 | 15.73 ± 0.16 |
| Carnitine (162.0 / 85.0) | 20.54 ± 0.3 | 19.3 ± 0.25 | 19.53 ± 0.17a | 19.6 ± 0.17 | 20.92 ± 0.17 | 20.27 ± 0.32b |
| cGMP (344.0 / 150.0) | 18.21 ± 0.41 | 17.5 ± 0.31 | 16.74 ± 0.52 | 17.1 ± 0.46 | 18.64 ± 0.65 | 18.00 ± 0.79 |
| Choline (104.0 / 60.0) | 24.16 ± 0.21 | 24.5 ± 0.22 | 24.1 ± 0.13 | 24.6 ± 0.18 | 24.59 ± 0.11 | 24.31 ± 0.15 |
| Citraconic Acid (129.0 / 85.0) | 20.57 ± 0.26 | 21.1 ± 0.15 | 20.36 ± 0.11 | 20.9 ± 0.15 | 21.29 ± 0.35 | 20.69 ± 0.17 |
| Citric Acid (191.0 / 111.0) | 18.71 ± 0.24 | 19.2 ± 0.32 | 19.08 ± 0.33 | 19.1 ± 0.17 | 19.20 ± 0.19 | 18.88 ± 0.22 |
| Citrulline (174.0 / 131.0) | 22.88 ± 0.4 | 24 ± 0.3 | 24.06 ± 0.46a | 23.6 ± 0.47 | 24.82 ± 0.27 | 23.99 ± 0.19b |
| Creatine (132.0 / 90.0) | 23.03 ± 0.11 | 21.2 ± 0.51 | 22.15 ± 0.26 | 22.5 ± 0.18 | 22.79 ± 0.24 | 22.54 ± 0.25 |
| Creatinine (114.0 / 44.0) | 20.54 ± 0.68 | 17.2 ± 0.22 | 18.24 ± 0.26a | 20 ± 1.09 | 18.11 ± 0.23 | 17.27 ± 0.36 |
| Cytidine (244.2 / 112.1) | 20.81 ± 0.18 | 21.2 ± 0.15 | 19.65 ± 0.26a | 20.3 ± 0.14b | 21.37 ± 0.37 | 20.74 ± 0.36 |
| Cytosine (112.0 / 95.0) | 17.52 ± 0.66 | 20.3 ± 0.44 | 18.09 ± 0.62 | 20.1 ± 0.74b | 17.29 ± 0.25 | 21.24 ± 0.48b |

| | | | | | | | |
|---|---|---|---|---|---|---|---|
| Deoxycarnitine (147.0 / 87.0) | 17.88 ± 0.42 | 17.5 ± 0.6 | 17.44 ± 0.47 | 16.9 ± 0.38 | | 19.81 ± 0.21 | 17.89 ± 0.44b |
| D-Leucic Acid (131.0 / 85.0) | 21.65 ± 0.28 | 22.1 ± 0.2 | 22.43 ± 0.5a | 22.9 ± 0.32 | | 20.51 ± 0.17 | 22.08 ± 0.21b |
| Fructose (179.0 / 89.0 (3)) | 18.67 ± 0.84 | 17.7 ± 0.33 | 18.44 ± 0.36 | 18.1 ± 0.75 | | 17.65 ± 0.35 | 18.29 ± 0.28 |
| Glucoronate (193.0 / 73.0) | 19.77 ± 0.5 | 19.6 ± 0.18 | 19.13 ± 0.32 | 20 ± 0.3b | | 19.39 ± 0.28 | 19.64 ± 0.26 |
| Glucosamine (180.1 / 162.0) | 16.85 ± 0.12 | 17.6 ± 0.11 | 16.68 ± 0.2 | 16.9 ± 0.4 | | 17.49 ± 0.24 | 17.51 ± 0.42 |
| Glucose (179.0 / 89.0) | 22.88 ± 0.57 | 22.5 ± 0.44 | 23.03 ± 0.05 | 23.7 ± 0.46 | | 21.99 ± 0.31 | 22.97 ± 0.16b |
| Glutamic acid (148.0 / 84.0) | 23.11 ± 0.22 | 23.7 ± 0.34 | 24.03 ± 0.29a | 23.1 ± 0.25 | | 25.22 ± 0.08 | 24.12 ± 0.25b |
| Glutamine (147.0 / 84.0) | 20.87 ± 0.36 | 20.2 ± 0.16 | 22.2 ± 0.2a | 21.5 ± 0.27 | | 23.16 ± 0.07 | 21.17 ± 0.18b |
| Glyceraldehyde (89.0 / 59.0) | 16.65 ± 0.43 | 16.7 ± 0.25 | 16.9 ± 0.15 | 17.5 ± 0.37 | | 15.99 ± 0.25 | 16.92 ± 0.18b |
| Glyceraldehyde (91.0 / 65.0) | 17.56 ± 0.22 | 17.6 ± 0.1 | 18.6 ± 0.16a | 17.9 ± 0.12b | | 19.42 ± 0.08 | 17.70 ± 0.08b |
| Glycerate (105.0 / 75.0) | 21.74 ± 0.26 | 20.9 ± 0.31 | 21 ± 0.17 | 20.3 ± 0.12b | | 21.26 ± 0.15 | 21.64 ± 0.16 |
| Glycine (76.0 / 30.1) | 15.21 ± 0.3 | 15.2 ± 0.26 | 15.77 ± 0.2 | 15.6 ± 0.36 | | 16.95 ± 0.09 | 15.67 ± 0.18b |
| Guanosine (284.2 / 152.1) | 19.89 ± 0.15 | 18.6 ± 0.39 | 16.93 ± 0.35a | 18.4 ± 0.7b | | 20.51 ± 0.36 | 17.83 ± 0.58b |
| Histamine (112.0 / 95.0 (2)) | 17.12 ± 0.75 | 16.5 ± 0.53 | 16.54 ± 0.17 | 16.9 ± 0.74 | | 18.03 ± 0.88 | 16.72 ± 0.45 |
| Histidine (156.0 / 110.0) | 21.72 ± 0.24 | 21.4 ± 0.17 | 22.46 ± 0.12a | 21.9 ± 0.2 | | 23.56 ± 0.12 | 21.58 ± 0.17b |
| Homoserine (120.0 / 74.0) | 20.05 ± 0.34 | 19.9 ± 0.17 | 21.04 ± 0.19a | 20.5 ± 0.3 | | 22.14 ± 0.06 | 20.58 ± 0.16b |
| Hydroxyproline (132.0 / 86.2) | 18.47 ± 0.15 | 18.4 ± 0.24 | 18.36 ± 0.1 | 18.8 ± 0.23 | | 18.24 ± 0.15 | 18.71 ± 0.22 |

| | | | | | | |
|---|---|---|---|---|---|---|
| Hypoxanthine (135.0 / 92.0) | 25.08 ± 0.3 | 26.2 ± 0.36 | 26.37 ± 0.28 | 26.1 ± 0.28 | 26.96 ± 0.21 | 26.52 ± 0.43 |
| Indole-3-Acetic Acid (174.0 / 130.0) | 17.1 ± 0.3 | 17.8 ± 0.24 | 16.65 ± 0.44 | 17.7 ± 0.13 | 15.98 ± 0.23 | 16.58 ± 0.22 |
| Inositol (179.0 / 87.0) | 19.68 ± 0.4 | 19.5 ± 0.22 | 19.43 ± 0.26 | 19.7 ± 0.15 | 19.55 ± 0.29 | 19.62 ± 0.26 |
| iso-Leucine (132.0 / 86.0 (2)) | 20.04 ± 0.35 | 19.4 ± 0.23 | 21.62 ± 0.17a | 20.3 ± 0.34b | 22.24 ± 0.09 | 20.16 ± 0.23b |
| Kynurenic Acid (188.0 / 144.0) | 20.2 ± 0.34 | 21.3 ± 0.2 | 20.2 ± 0.26 | 21.1 ± 0.26b | 18.92 ± 0.19 | 19.70 ± 0.2b |
| lactate (89.0 / 43.0) | 22.14 ± 0.41 | 21.3 ± 0.63 | 21.85 ± 0.38 | 22.8 ± 0.36 | 21.20 ± 0.24 | 21.59 ± 0.26 |
| Lactose (341.0 / 59.0) | 19.46 ± 0.42 | 20.1 ± 0.71 | 19.25 ± 0.22 | 20.8 ± 0.75 | 18.37 ± 0.07 | 19.44 ± 0.24 |
| Leucine (132.0 / 86.0) | 22.36 ± 0.26 | 21.4 ± 0.19 | 23.66 ± 0.14a | 22.6 ± 0.27b | 24.45 ± 0.07 | 22.47 ± 0.18b |
| Linoleic Acid (279.1 / 261.0) | 21.82 ± 0.44 | 22.5 ± 0.19 | 23.16 ± 0.17a | 23.1 ± 0.06 | 21.03 ± 0.47 | 22.46 ± 0.28b |
| Linolenic Acid (277.1 / 259.0) | 18.29 ± 0.46 | 18.8 ± 0.23 | 19.69 ± 0.2b | 19.7 ± 0.08 | 17.29 ± 0.29 | 18.83 ± 0.39b |
| Lysine (147.0 / 84.0 (2)) | 22.41 ± 0.28 | 22.8 ± 0.22 | 23.97 ± 0.24a | 22.7 ± 0.27b | 24.77 ± 0.07 | 22.78 ± 0.18b |
| Malate (133.0 / 115.0) | 22.03 ± 0.35 | 22.3 ± 0.38 | 21.82 ± 0.12 | 21.1 ± 0.21 | 22.12 ± 0.12 | 22.88 ± 0.61 |
| Maleic Acid (115.0 / 71.0 (2)) | 20.32 ± 0.3 | 21.2 ± 0.14 | 20.37 ± 0.18 | 21 ± 0.12 | 21.07 ± 0.14 | 20.67 ± 0.16 |
| Malondialdehyde (71.0 / 41.0) | 18.75 ± 0.55 | 18.8 ± 0.36 | 18.82 ± 0.24 | 19.6 ± 0.46 | 18.06 ± 0.26 | 18.98 ± 0.16b |
| Margaric Acid (269.1 / 251.3) | 18.85 ± 0.11 | 19.4 ± 0.28 | 19.63 ± 0.34 | 19.3 ± 0.06 | 18.95 ± 0.16 | 18.31 ± 0.29 |
| Methionine (150.0 / 61.0) | 18.76 ± 0.29 | 18.5 ± 0.21 | 20.25 ± 0.29a | 19.3 ± 0.31 | 21.47 ± 0.08 | 19.18 ± 0.21b |
| N-AcetylGlycine (116.0 / 74.0) | 16.5 ± 0.75 | 18.7 ± 0.49 | 15.96 ± 0.08 | 17.8 ± 0.62 | 15.55 ± 0.11 | 19.33 ± 0.19b |
| N-Acetylneuraminate (308.1 / 87.0) | 24.47 ± 0.44 | 24.3 ± 0.32 | 24.01 ± 0.21a | 24.5 ± 0.31 | 24.89 ± 0.12 | 24.79 ± 0.16 |

| | | | | | | |
|---|---|---|---|---|---|---|
| Nicotinic Acid (122.0 / 78.0) | 23.04 ± 0.18 | 24.6 ± 0.35 | 23.64 ± 0.29 | 23.6 ± 0.18 | 23.97 ± 0.17 | 23.88 ± 0.34 |
| Ornithine (133.0 / 70.0) | 17.79 ± 0.25 | 19.1 ± 0.29 | 18.97 ± 0.17a | 18.1 ± 0.37b | 18.44 ± 0.12 | 18.34 ± 0.21 |
| Orotate (155.0 / 111.0) | 19.57 ± 0.67 | 19.9 ± 0.37 | 19.4 ± 0.77 | 19.8 ± 0.35 | 18.68 ± 0.54 | 19.78 ± 0.34 |
| Oxalacetate (131.0 / 113.0) | 19.94 ± 0.39 | 18.4 ± 0.1 | 21.6 ± 0.17a | 19.1 ± 0.17b | 22.60 ± 0.06 | 19.87 ± 0.30b |
| Pentothenate (218.1 / 88.0) | 22.14 ± 0.37 | 24.2 ± 0.43 | 22.66 ± 0.44 | 22.1 ± 0.33 | 22.89 ± 0.23 | 23.04 ± 0.37 |
| Phenylalanine (166.0 / 120.0) | 23.45 ± 0.26 | 22.7 ± 0.19 | 24.71 ± 0.17a | 23.7 ± 0.21b | 25.55 ± 0.08 | 23.48 ± 0.18b |
| Pipecolate (130.0 / 84.0) | 17.61 ± 0.42 | 18.9 ± 0.41 | 17.25 ± 0.41 | 16.7 ± 0.17 | 17.79 ± 0.18 | 18.24 ± 0.41 |
| PPA (163.0 / 91.0) | 17 ± 0.17 | 16.8 ± 0.23 | 17.24 ± 0.15 | 17.4 ± 0.31 | 18.29 ± 0.14 | 18.38 ± 0.40 |
| Proline (116.0 / 70.0) | 20.04 ± 0.34 | 20.8 ± 0.32 | 20.76 ± 0.21a | 20.9 ± 0.29b | 21.50 ± 0.24 | 20.94 ± 0.14 |
| Pyroglutamic Acid (130.0 / 83.9) | 16.3 ± 0.36 | 17.3 ± 0.32 | 16.24 ± 0.24 | 16.5 ± 0.21 | 17.04 ± 0.20 | 16.87 ± 0.25 |
| Pyruvate (87.0 / 43.0) | 17.17 ± 0.1 | 17.7 ± 0.16 | 17.4 ± 0.24 | 18 ± 0.29b | 18.67 ± 0.09 | 18.24 ± 0.33 |
| Quinolinic Acid (168.0 / 150.0) | 17.52 ± 0.28 | 18.3 ± 0.24 | 17.66 ± 0.14 | 17.7 ± 0.13 | 17.59 ± 0.17 | 17.50 ± 0.19 |
| Reduced glutathione (306.3 / 143.1) | 15.41 ± 0.3 | 15.2 ± 0.17 | 17 ± 0.26a | 15.9 ± 0.21b | 17.54 ± 0.20 | 15.10 ± 0.19b |
| Serine (106.0 / 60.0 (2)) | 20.11 ± 0.28 | 19.6 ± 0.11 | 21.03 ± 0.17a | 20.3 ± 0.25 | 22.05 ± 0.04 | 20.45 ± 0.16b |
| Sorbitol (181.0 / 89.0) | 18.3 ± 0.52 | 18.2 ± 0.3 | 18.02 ± 0.35 | 17.8 ± 0.48 | 17.69 ± 0.10 | 18.10 ± 0.22 |
| Succinate (117.0 / 73.0) | 22.66 ± 0.4 | 21.4 ± 0.43 | 20.93 ± 0.31a | 22.3 ± 0.34b | 20.25 ± 0.20 | 21.92 ± 0.21b |
| Sucrose (341.0 / 59.0 (2)) | 19.13 ± 0.9 | 18.8 ± 0.56 | 18.04 ± 0.16a | 18.5 ± 0.68 | 17.74 ± 0.22 | 17.65 ± 0.19 |
| Taurine (126.0 / 108.0) | 20.23 ± 0.68 | 18.6 ± 0.43 | 20.14 ± 0.32 | 19 ± 0.15b | 21.64 ± 0.25 | 21.16 ± 0.31 |
| Taurocholate (514.5 / 124.0) | 18.01 ± 0.5 | 16.7 ± 0.29 | 17.8 ± 0.44 | 17.5 ± 0.25 | 20.09 ± 0.58 | 18.30 ± 0.14b |

| | | | | | | | |
|---|---|---|---|---|---|---|---|
| Threonine (120.0 / 74.0 (2)) | 20.06 ± 0.33 | 19.9 ± 0.18 | 21.05 ± 0.19a | 20.5 ± 0.31 | | 22.13 ± 0.07 | 20.56 ± 0.15b |
| Tryptophan (205.1 / 146.0) | 19.84 ± 0.32 | 19.5 ± 0.38 | 20.82 ± 0.15a | 19.9 ± 0.21 | | 21.92 ± 0.06 | 19.74 ± 0.19b |
| Tyramine (138.0 / 121.0) | 21.03 ± 0.27 | 21.8 ± 0.27 | 21.04 ± 0.09 | 21.3 ± 0.08 | | 21.21 ± 0.16 | 21.04 ± 0.22 |
| Tyrosine (182.1 / 136.0) | 20.88 ± 0.25 | 20.7 ± 0.1 | 22.12 ± 0.18a | 21.4 ± 0.18b | | 22.99 ± 0.07 | 21.03 ± 0.12b |
| Uracil (111.0 / 42.0) | 23.54 ± 0.23 | 24.7 ± 0.31 | 23.46 ± 0.34 | 24 ± 0.44b | | 24.04 ± 0.23 | 24.87 ± 0.27b |
| Urate (167.0 / 124.0) | 18.65 ± 0.86 | 19.6 ± 0.58 | 16.85 ± 0.08 | 18.5 ± 0.57b | | 18.61 ± 0.33 | 19.23 ± 0.16 |
| Uridine (245.2 / 113.1) | 19.17 ± 0.27 | 18.7 ± 0.21 | 18.4 ± 0.2 | 18.6 ± 0.33 | | 19.50 ± 0.34 | 18.00 ± 0.31b |
| Valine (118.0 / 72.0) | 18.57 ± 0.28 | 18.3 ± 0.23 | 19.86 ± 0.16a | 18.9 ± 0.27b | | 20.75 ± 0.05 | 18.84 ± 0.17b |
| Xanthine (151.0 / 108.0) | 24.76 ± 0.27 | 25.2 ± 0.42 | 24.41 ± 0.44 | 25.2 ± 0.51b | | 24.76 ± 0.24 | 26.07 ± 0.30b |
| Xanthurenic Acid (204.1 / 160.0) | 19.59 ± 0.44 | 19.6 ± 0.28 | 19.69 ± 0.24 | 18.9 ± 0.17b | | 19.65 ± 0.23 | 19.25 ± 0.18 |

Each value represents the mean ± SEM of normalized and imputed abundance data of metabolites generated from 6 mice per group. Significant differences between groups are shown for an FDR <0.05: a = significant difference between adult AL and old AL, b = significant difference between old AL and old CR. The numbers in parenthesis by the metabolite names indicates Parts/million (PPM)/Multiple Reaction Monitoring (MRM).

**Table 5S. List of the top 20 Transcripts that Show a Significant and 2-Fold Change [P (Corr) <=0.05) with Age or CR in C57BL/6 Mice**

| Ensembl ID | Entrez ID | Log FC | FC (abs) | Regulation | Gene Symbol |
|---|---|---|---|---|---|
| **Transcripts that change with age (Old AL vs Adult AL) in C57BL/6 mice** | | | | | |
| ENSMUSG00000079260 | 100504715 | 5.939521 | 61.37254 | up | Tmppe |
| ENSMUSG00000020826 | 18126 | 5.692607 | 51.71844 | up | Nos2 |
| ENSMUSG00000079494 | 69049 | 4.578009 | 23.8846 | up | Nat8f5 |
| ENSMUSG00000030004 | 68396 | 4.480648 | 22.32593 | up | Nat8 |
| ENSMUSG00000044988 | 83428 | 4.337706 | 20.21992 | up | Ucn3 |
| ENSMUSG00000051262 | 93674 | 4.168231 | 17.97888 | up | Nat8f3 |
| ENSMUSG00000035394 | 74453 | 3.714172 | 13.12433 | up | Cfap53 |
| ENSMUSG00000025194 | 100038628 | 3.380169 | 10.41195 | up | Gm10768 |
| ENSMUSG00000087361 | 68400 | 3.302278 | 9.864719 | up | 0610043K17Rik |
| ENSMUSG00000031302 | 245537 | 3.136771 | 8.795529 | up | Nlgn3 |
| ENSMUSG00000002265 | 18616 | -4.84128 | 28.66614 | down | Peg3 |
| ENSMUSG00000054423 | 27062 | -4.61523 | 24.5088 | down | Cadps |
| ENSMUSG00000055193 | 317652 | -4.57431 | 23.82348 | down | Klk15 |
| ENSMUSG00000025991 | 227231 | -4.55286 | 23.47186 | down | Cps1 |
| ENSMUSG00000031430 | 78789 | -3.95537 | 15.51261 | down | Vsig1 |
| ENSMUSG00000055567 | 329178 | -3.80721 | 13.99862 | down | Unc80 |
| ENSMUSG00000027674 | 58869 | -3.75772 | 13.52654 | down | Pex5l |
| ENSMUSG00000031292 | 382253 | -3.7395 | 13.35675 | down | Cdkl5 |
| ENSMUSG00000074796 | 269356 | -3.70428 | 13.03465 | down | Slc4a11 |
| ENSMUSG00000009378 | 240638 | -3.61335 | 12.23844 | down | Slc16a12 |
| **Transcripts that change with CR (Adult CR vs Adult AL) in C57BL/6 mice** | | | | | |
| ENSMUSG00000020826 | 18126 | 2.707298 | 6.530971 | up | Nos2 |
| ENSMUSG00000013611 | 66696 | 2.399487 | 5.276156 | up | Snx31 |
| ENSMUSG00000031538 | 18791 | 2.322075 | 5.00051 | up | Plat |
| ENSMUSG00000061615 | 319172 | 2.292758 | 4.899918 | up | Hist1h2ab |
| ENSMUSG00000033676 | 14402 | 2.282655 | 4.865725 | up | Gabrb3 |

| | | | | | |
|---|---|---|---|---|---|
| ENSMUSG00000044988 | 83428 | 2.013193 | 4.036746 | up | Ucn3 |
| ENSMUSG00000108218 | 257871 | 1.958138 | 3.8856 | up | Olfr1372-ps1 |
| ENSMUSG00000015702 | 71790 | 1.919249 | 3.782262 | up | Anxa9 |
| ENSMUSG00000114442 | 100038627 | 1.817609 | 3.524965 | up | F630042J09Rik |
| ENSMUSG00000051262 | 93674 | 1.812597 | 3.512741 | up | Nat8f3 |
| ENSMUSG00000074796 | 269356 | -3.10644 | 8.612539 | down | Slc4a11 |
| ENSMUSG00000055193 | 317652 | -2.47794 | 5.571014 | down | Klk15 |
| ENSMUSG00000043165 | 16939 | -2.25796 | 4.783151 | down | Lor |
| ENSMUSG00000030205 | 93746 | -2.1144 | 4.330094 | down | Gprc5d |
| ENSMUSG00000031430 | 78789 | -1.92479 | 3.796819 | down | Vsig1 |
| ENSMUSG00000099003 | 102465626 | -1.90968 | 3.757266 | down | Mir7035 |
| ENSMUSG00000022225 | 17228 | -1.87459 | 3.666969 | down | Cma1 |
| ENSMUSG00000044083 | 100504221 | -1.84962 | 3.604044 | down | Efcab8 |
| ENSMUSG00000064925 | 104433 | -1.81453 | 3.517459 | down | Snora62 |
| | 99169 | -1.7873 | 3.451673 | down | AU015228 |
| **Transcripts that change with CR (Old CR vs Old AL) in old C57BL/6 mice** | | | | | |
| ENSMUSG00000027068 | 241452 | 3.636485 | 12.43629 | up | Dhrs9 |
| ENSMUSG00000007682 | 13371 | 3.547132 | 11.68942 | up | Dio2 |
| ENSMUSG00000030963 | 22242 | 3.267541 | 9.630031 | up | Umod |
| ENSMUSG00000030954 | 67133 | 3.209645 | 9.251227 | up | Gp2 |
| ENSMUSG00000078302 | 15229 | 3.118569 | 8.685258 | up | Foxd1 |
| ENSMUSG00000074796 | 269356 | 3.05619 | 8.317731 | up | Slc4a11 |
| ENSMUSG00000025991 | 227231 | 2.695655 | 6.478479 | up | Cps1 |
| ENSMUSG00000030205 | 93746 | 2.538617 | 5.810316 | up | Gprc5d |
| ENSMUSG00000024912 | 14283 | 2.5329 | 5.78734 | up | Fosl1 |
| ENSMUSG00000056716 | 432436 | 2.498507 | 5.651002 | up | Gm5420 |
| ENSMUSG00000079260 | 100504715 | -4.44553 | 21.789 | down | Tmppe |
| ENSMUSG00000020826 | 18126 | -4.24985 | 19.02536 | down | Nos2 |
| ENSMUSG00000030483 | 13088 | -3.38368 | 10.43735 | down | Cyp2b10 |
| ENSMUSG00000039787 | 99151 | -3.02324 | 8.129922 | down | Cercam |

| | | | | | |
|---|---|---|---|---|---|
| ENSMUSG00000027209 | 75823 | -2.65985 | 6.319671 | down | Fam227b |
| ENSMUSG00000038550 | 229599 | -2.57473 | 5.957578 | down | Ciart |
| ENSMUSG00000000489 | 18591 | -2.528 | 5.767729 | down | Pdgfb |
| ENSMUSG00000044988 | 83428 | -2.51122 | 5.701023 | down | Ucn3 |
| ENSMUSG00000062017 | 67928 | -2.49858 | 5.651283 | down | Abca14 |
| ENSMUSG00000063171 | 66184 | -2.45845 | 5.496249 | down | Rps4l |

The top 20 transcripts that show a significant fold change (FC) with either age or CR in adult and old C57BL/6 mice. Log Fold Change (Log FC) between two conditions is the difference between their respective average normalized signal values. Absolute FC (FC (abs)) is computed as (sign of Log FC) ×2|log FC|.
